# De novo mapping of the apicomplexan Ca^2+^-responsive proteome

**DOI:** 10.1101/2022.05.25.493445

**Authors:** Alice L. Herneisen, Zhu-Hong Li, Alex W. Chan, Silvia NJ Moreno, Sebastian Lourido

## Abstract

Apicomplexan parasites cause persistent mortality and morbidity worldwide through diseases including malaria, toxoplasmosis, and cryptosporidiosis. Ca^2+^ signaling pathways have been repurposed in these eukaryotic pathogens to regulate parasite-specific cellular processes governing the transition between the replicative and lytic phases of the infectious cycle. Despite the presence of conserved Ca^2+^-responsive proteins, little is known about how specific signaling elements interact to impact pathogenesis. We mapped the Ca^2+^-responsive proteome of the model apicomplexan *T. gondii* via time-resolved phosphoproteomics and thermal proteome profiling. The waves of phosphoregulation following PKG activation and stimulated Ca^2+^ release corroborate known physiological changes but identify specific proteins operating in these pathways. Thermal profiling of parasite extracts identified many expected Ca^2+^-responsive proteins, such as parasite Ca^2+^-dependent protein kinases. Our approach also identified numerous Ca^2+^-responsive proteins that are not predicted to bind Ca^2+^, yet are critical components of the parasite signaling network. We characterized protein phosphatase 1 (PP_1_) as a Ca^2+^-responsive enzyme that relocalized to the parasite apex upon Ca^2+^ store release. Conditional depletion of PP_1_ revealed that the phosphatase regulates Ca^2+^ uptake to promote parasite motility. PP_1_ may thus be partly responsible for Ca^2+^-regulated serine/threonine phosphatase activity in apicomplexan parasites.

## INTRODUCTION

Apicomplexan parasites cause persistent mortality and morbidity worldwide through diseases including malaria, toxoplasmosis, and cryptosporidiosis (Havelaar et al., 2015). The phylum member *Toxoplasma gondii* alone infects >2 billion people. As obligate intracellular pathogens, apicomplexans are exquisitely tuned to transduce environmental signals into programs of motility, replication, and quiescence responsible for parasite pathogenesis and spread (Bisio and Soldati-favre, 2019). Ca^2+^ signals and their downstream effectors are a part of the signaling cascade that triggers changes impacting almost every cellular function (Lourido and Moreno, 2015). Signaling begins with release of Ca^2+^ from intracellular stores or influx through plasma membrane channels, resulting in diverse downstream events central to parasite virulence, including secretion of adhesive proteins, motility, and invasion into and egress from host cells. Together, these cellular processes orchestrate a dramatic transition from the replicative to the kinetic phase of the life cycle that allows parasites to spread to new host cells. Signal-transducing components downstream of Ca^2+^ release are largely unknown yet likely essential for apicomplexan viability and virulence (Lourido and Moreno, 2015; Nagamune and Sibley, 2006).

Ca^2+^ can change a protein’s state by direct binding or indirect effects such as triggering post-translational modification (PTM), interaction with other proteins, or relocalization. Indirect effects are fundamental to the propagation and amplification of signals across the Ca^2+^-regulated network, and in most organisms they are largely mediated by three classes of Ca^2+^-binding proteins: Ca^2+^-regulated kinases, Ca^2+^-regulated phosphatases, and calmodulin (CaM) and related proteins (Villalobo et al., 2019). Genomic searches for canonical Ca^2+^-binding domains in apicomplexans have identified several individual proteins involved in transducing and effectuating Ca^2+^ signals (Farrell et al., 2012; Huet et al., 2018; W. Li et al., 2021; McCoy et al., 2017), like kinases, phosphatases, and transporters (Hortua Triana et al., 2018; Lourido et al., 2012, 2010; Luo et al., 2005; Márquez-Nogueras et al., 2021). However, many of the key signaling elements involved in fundamental Ca^2+^ responses—including the channels responsible for its stimulated release—are either missing from apicomplexan genomes or have diverged beyond recognition, suggesting that eukaryotic pathogens evolved novel pathways for Ca^2+^ mobilization and transduction (Billker et al., 2009; Lourido and Moreno, 2015).

In apicomplexans, Ca^2+^-dependent protein kinases (CDPKs) have garnered the most attention as the only known Ca^2+^-regulated kinases in the phylum (Billker et al., 2009). These kinases possess intrinsic Ca^2+^-binding sites and do not rely on CaM-like mammalian CaMKs. In *T. gondii*, several of these CDPKs trigger parasite motility (Lourido et al., 2012, 2010; McCoy et al., 2012; Smith et al., 2022; Treeck et al., 2014). Within the parasite Ca^2+^ signaling field, dephosphorylation has garnered comparatively little attention (Yang and Arrizabalaga, 2017). The roles of the prototypical CaM and the Ca^2+^/CaM-dependent phosphatase calcineurin have only been phenotypically examined in parasites, and their client proteins remain largely unknown (Paul et al., 2015; Philip and Waters, 2015). Although key players are conserved and essential across the Apicomplexa, no systematic efforts have been undertaken to globally map the Ca^2+^ signaling pathways of these pathogens.

Ca^2+^ signaling pathways have been repurposed in apicomplexans to regulate parasite-specific cellular processes governing the transition between the replicative and kinetic phases of the infectious cycle (Brown et al., 2020; Pace et al., 2020). Despite the presence of conserved Ca^2+^-responsive proteins (Lourido and Moreno, 2015; Nagamune and Sibley, 2006), uncovering the Ca^2+^ signaling architecture of apicomplexans demands a reevaluation of the entire network to understand how specific signaling elements interact to impact pathogenesis. We present an atlas of Ca^2+^-regulated proteins in the model apicomplexan *T. gondii*, assembled from high-dimensional proteomic datasets. The physiological changes associated with stimulated motility in the asexual stages of parasites have been characterized for decades (Blader et al., 2015). Our approach identified at once hundreds of molecular components underpinning these processes. We find numerous Ca^2+^-responsive proteins that are not predicted to bind Ca^2+^, yet operate at critical junctures in the parasite signaling network. From this analysis, the protein phosphatase PP_1_ emerges as an unanticipated Ca^2+^-responsive phosphatase.

## RESULTS

### Sub-minute phosphoproteomics reveals the topology of Ca^2+^-dependent signaling processes

Parasite Ca^2+^ fluxes can arise from different sources (Bisio et al., 2019; Bisio and Soldati-favre, 2019; Lourido et al., 2012). Ca^2+^ release from intracellular stores is mobilized by cyclic nucleotide signaling (Brown et al., 2016; Sidik et al., 2016b). Ca^2+^ entry from the extracellular space has also been observed (Pace et al., 2014; Vella et al., 2021). We can emulate these endogenous signaling pathways by treating isolated parasites with zaprinast, which stimulates PKG activation by inhibiting phosphodiesterases that degrade cGMP (Brown et al., 2016; Lourido et al., 2012). Zaprinast-stimulated motility occurs rapidly and in defined sequence in apicomplexans: initially, an increase in cGMP activates parasite protein kinase G (PKG), which phosphorylates substrates and stimulates Ca^2+^ release from internal stores (Brown et al., 2020; Lourido and Moreno, 2015). Ca^2+^-dependent protein kinases, such as *Tg*CDPK1 and *Tg*CDPK3, synergize with PKG to effectuate microneme secretion and parasite motility (Brown et al., 2016; Lourido et al., 2012, 2010; McCoy et al., 2012). This signaling cascade is active within seconds of cGMP elevation; however, existing *T. gondii* phosphoproteomes compare changes at a single time point following Ca^2+^-ionophore stimulation (Treeck et al., 2014), providing only a snapshot of diverging signaling states. Here, we add kinetic resolution to these signaling pathways. We quantified dynamic changes in the phosphoproteome within a minute of stimulation with zaprinast and thus activation of the cGMP/Ca^2+^ pathway.

We collected five timepoints in the 60 seconds following stimulation (0, 5, 10, 30, and 60 s), as well as three DMSO-treated matched timepoints (0, 10, and 30 s), in biological duplicate (**Figure 1A**). Using TMTpro labeling methods (Li et al., 2020), we multiplexed 16 samples, allowing us to analyze a complete time course, with replicates and controls, in a single MS experiment. Phosphopeptides were enriched from the rest of the sample using sequential metal-oxide affinity chromatography, which maximizes phosphopeptide capture (Tsai et al.,2014). Our experiments quantified 4,055 parasite proteins, none of which exhibited more than a two-fold change in abundance in the 60 seconds following stimulation (**Figure 1 Supplement**).

**Figure 1.**
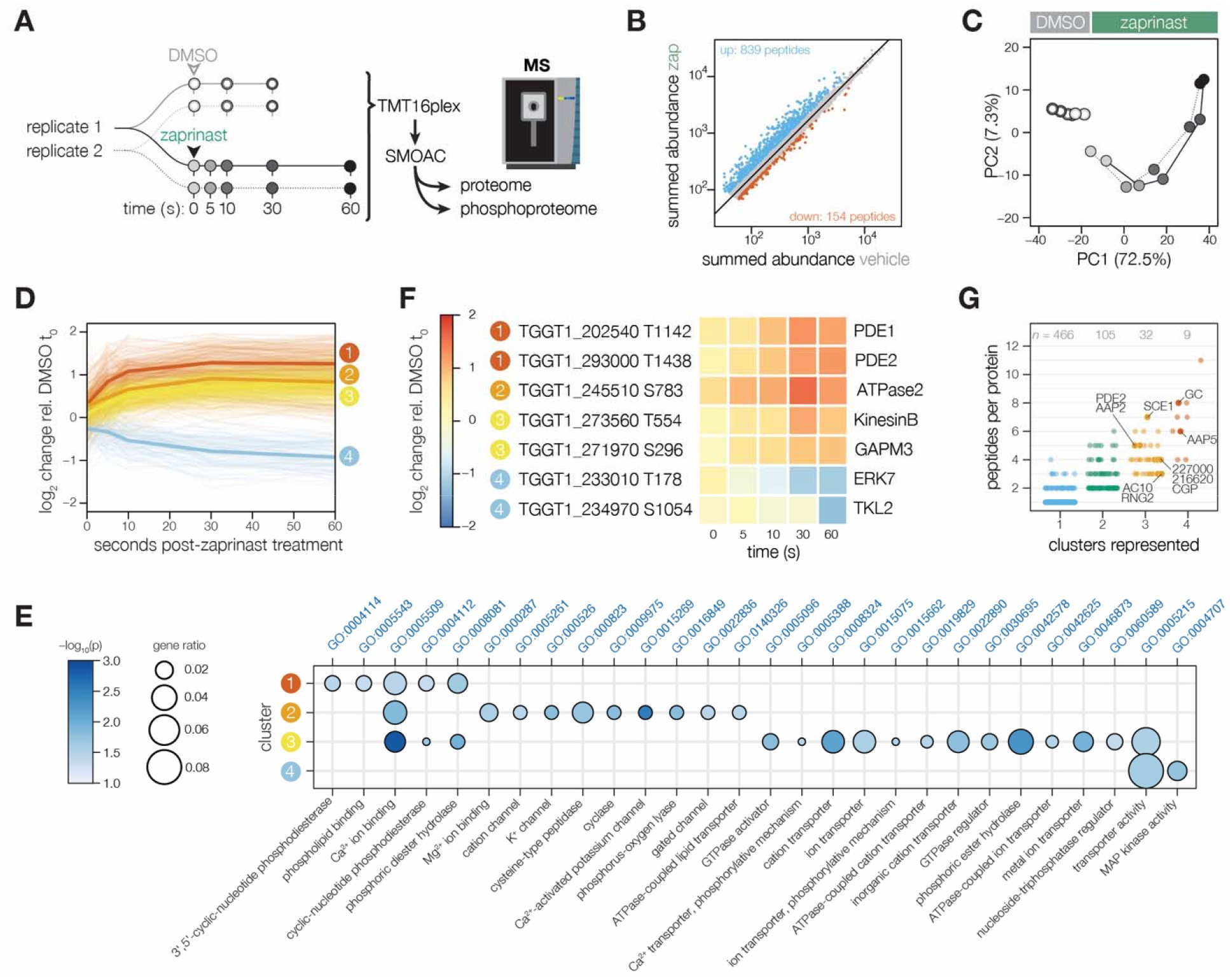
Phosphoregulation triggered by Ca^2+^ release. **(A)** Schematic of the sub-minute phosphoproteomics experiments with the Ca^2+^ signaling agonist zaprinast. **(B)** The summed abundances of unique phosphopeptides during zaprinast or vehicle (DMSO) treatment. The abundance ratios were transformed into a modified Z-score and were used to threshold increasing (Z > 3; *blue*) or decreasing (Z < -1.8; *orange)* phosphopeptides. **(C)** Principal component analysis of phosphopeptides identified as significantly changing. Symbols follow the schematic in **A. (D)** Gaussian mixture-model-based clustering of phosphopeptides changing during zaprinast treatment. Solid lines show the median relative abundance of each cluster. Opaque lines show the individual phosphopeptides belonging to each cluster. **(E)** GO terms enriched among phosphopeptides changing with zaprinast treatment, grouped by cluster. Gene ratio is the proportion of proteins with the indicated GO term to the total number of proteins belonging to each cluster. Significance was determined with a hypergeometric test; only GO terms with *p* < 0.05 are shown. Redundant GO terms were removed. **(F)** Examples of phosphopeptides belonging to each cluster. **(G)** The number of clusters each phosphoprotein belongs to, plotted against the number of changing phosphopeptides belonging to each protein. Gene names or IDs indicate proteins discussed in the text.

Given the paucity of known phosphoreglatory interactions in apicomplexans compared to other organisms (Weiss et al., 2020), we employed several analysis approaches to maximize the identification of changing phosphosites. We first calculated phosphoregulation scores by summing peptide abundances of vehicle (DMSO) and zaprinast-treated samples, taking their ratios, and standardizing the values with a modified Z score (**Figure 1B**). From a phosphoproteome of 11,755 unique peptides with quantification values (belonging to 2,690 phosphoproteins), 839 phosphopeptides increased in abundance three modified Z-scores above the median, whereas 154 decreased 1.8 Z-scores below the median. Principal component analysis on the significant peptides distinguished the agonist treatment and time course kinetics in the two principal components accounting for the greatest variability in the data (**Figure 1C** and **Figure 1 Supplement**).

### Kinetically resolved clusters reveal regulatory subnetworks during zaprinast stimulation

We leveraged the kinetic resolution of our comprehensive phosphoproteomics datasets to identify subregulatory networks. A Gaussian mixture-model clustering algorithm (Invergo et al., 2017) heuristically resolved four clusters for phosphopeptides arising from zaprinast treatment: three clusters increasing with different kinetics, and one decreasing (**Figure 1D**). On average, the 173 phosphopeptides belonging to cluster 1 increased sharply in abundance within 5 seconds of treatment and continued to increase in the remainder of the time course (**Figure 1D**), suggesting that they belonged to the first wave of phosphoregulation. This cluster was enriched for phosphoproteins associated with phosphodiesterase activity, phospholipid binding, and Ca^2+^ binding (**Figure 1E**), including PDE1, PDE2, PI4,5K, PI3,4K, PI-PLC, a phosphatidylinositol-3,4,5-triphosphate 5-phosphatase, a putative Sec14, *Tg*CDPK2A and *Tg*CDPK7, and PPM2B (**Table 1** and **Figure 1F**).

**Table 1.**
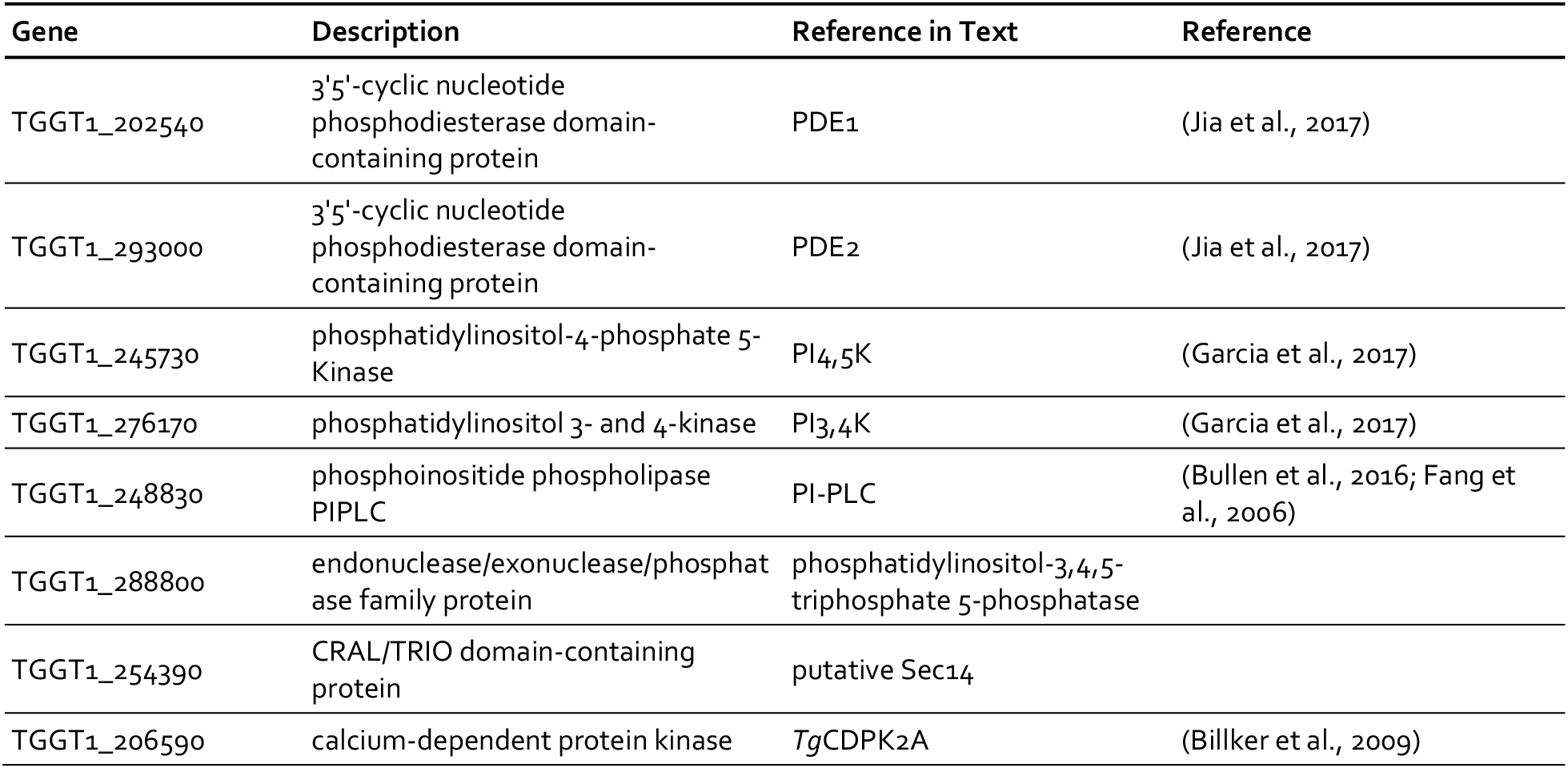
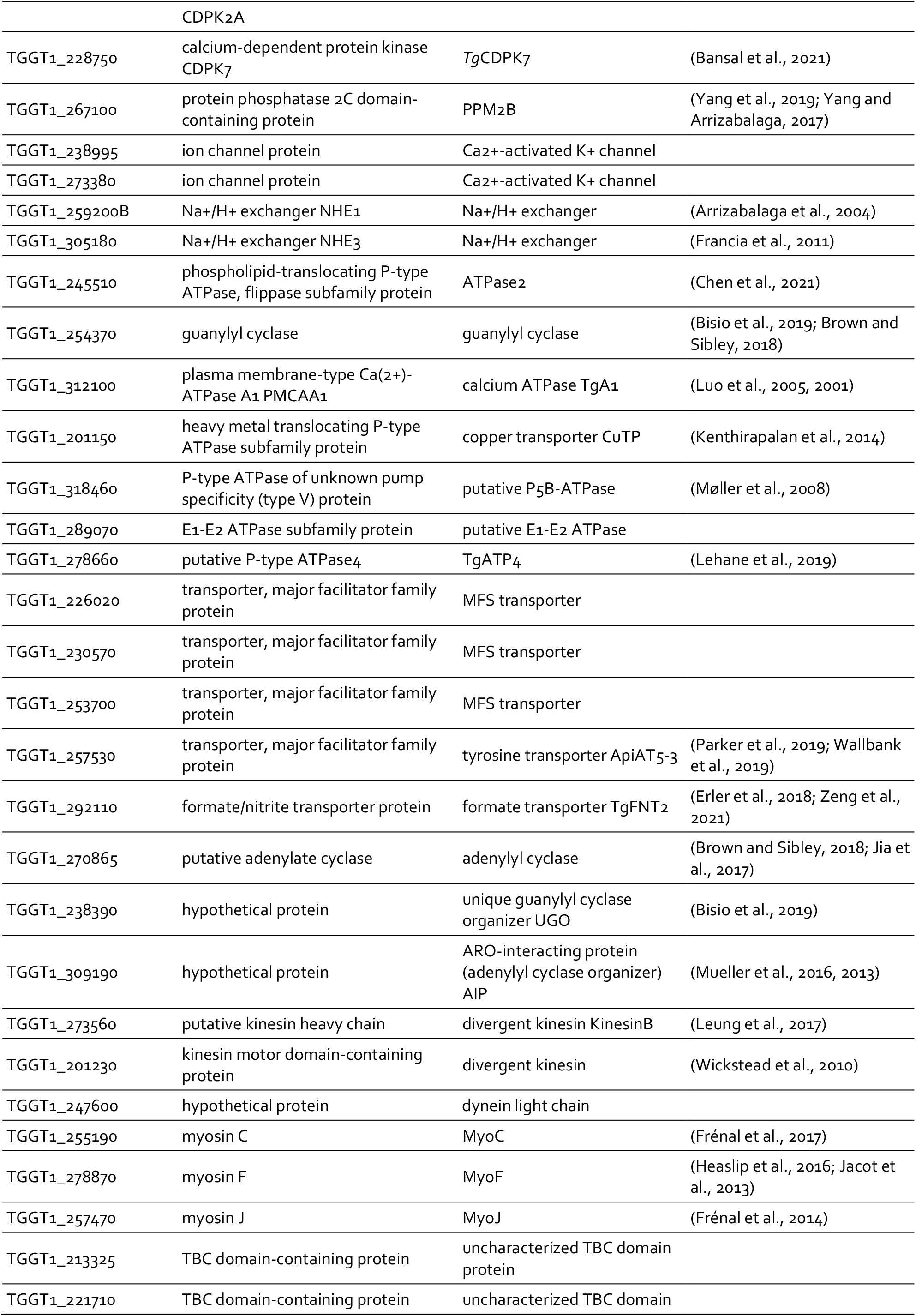
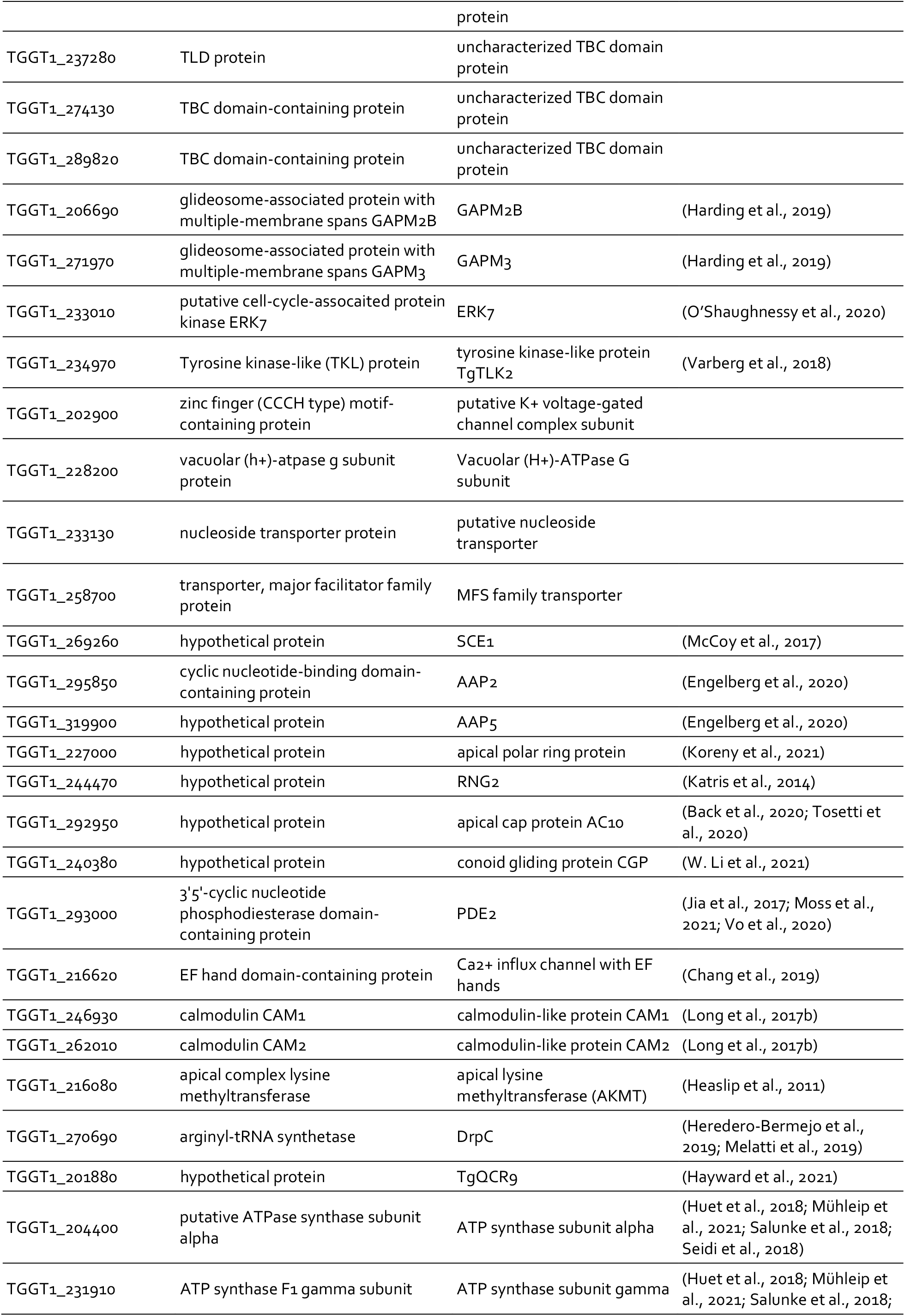
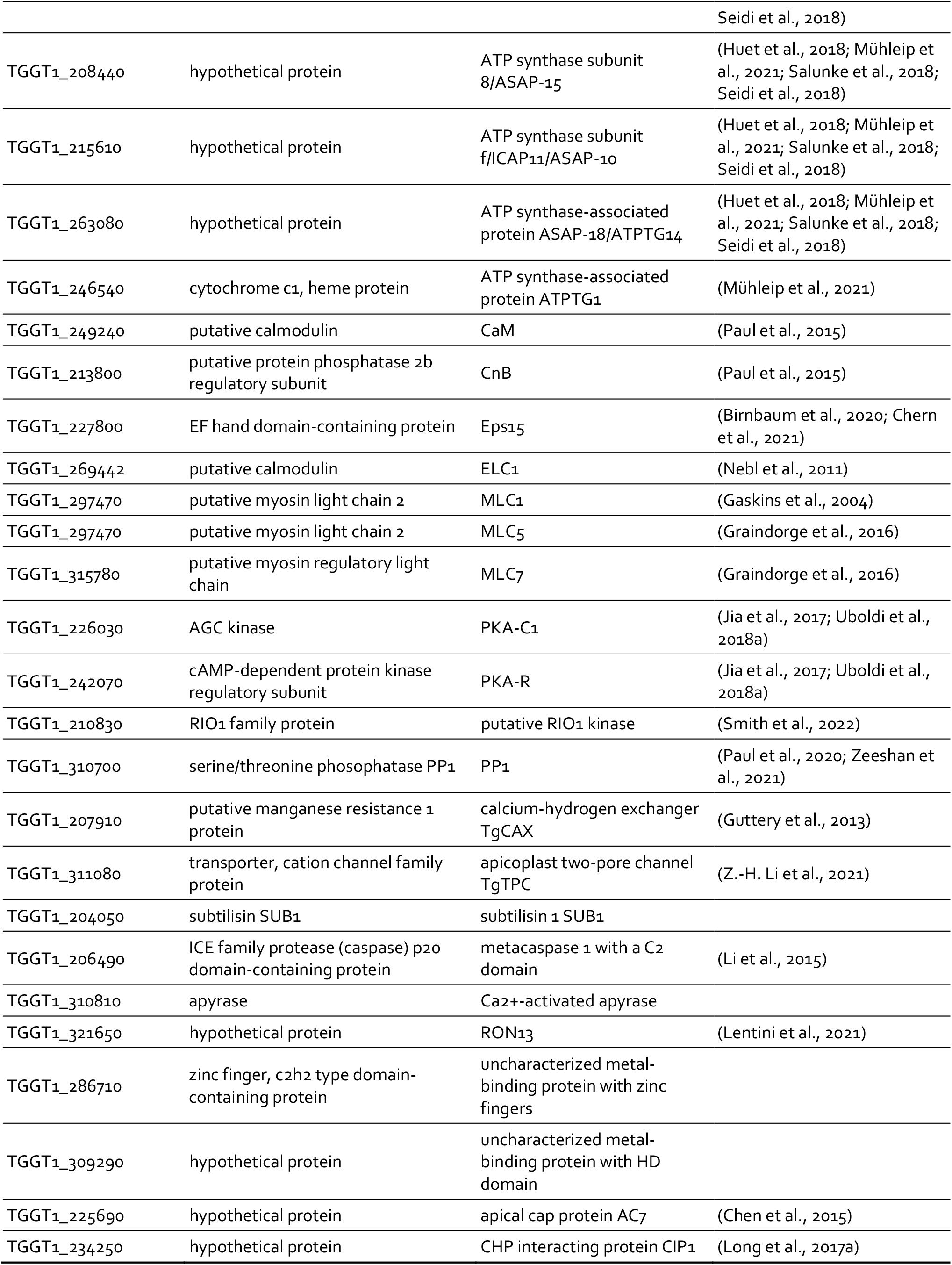
Gene IDs of proteins discussed in the text.

Peptides in clusters 2 and 3 (173 and 527, respectively) increased more gradually and exhibited lower fold-changes than cluster 1 sites (**Figure 1D**). Cluster 2 was notably enriched in proteins functioning in transport of monovalent ions and lipids, cyclase activity, and Ca^2+^ binding (**Figure 1E**). This set includes two putative Ca^2+^-activated K^+^ channels and the sodium-hydrogen exchangers NHE1 and NHE3 (**Table 1**). ATPase2 and the guanylyl cyclase in this cluster both have putative phospholipid-transporting ATPase domains. PI4,5K, PI3,4K, and *Tg*CDPK2A, noted in cluster 1, have additional phosphoregulatory sites belonging to cluster 2. Clusters 2 and 3 are enriched for phosphoproteins with metal transporter activity, including ATPases such as *Tg*A1, the putative copper transporter CuTP, a putative P5B-ATPase, a putative E1-E2 ATPase, and *Tg*ATP4. As observed in other *T. gondii* Ca^2+^-stimulated phosphoproteomes, at later time points small-molecule transporters are phosphorylated, including MFS transporters, ApiAT5-3, and *Tg*FNT2. The guanylyl and adenylyl cyclases are also extensively modified, as are the cyclase organizers UGO and AIP.

Cluster 3, which represents a latter wave of phosphoregulation, is uniquely enriched in proteins functioning in subcellular remodeling, vesicle trafficking, and glideosome activity (**Figure 1E**). This set includes two divergent kinesins (KinesinB and TGGT1_201230), a dynein light chain, and the myosin motors MyoC, MyoF, and MyoJ (**Table 1**). Several uncharacterized GTPase regulators are phosphorylated later in the zaprinast response, including putative ARF1 activators (TGGT1_225310 and TGGT1_266830) as well as five uncharacterized TBC domain proteins. The glideosome-associated membrane proteins GAPM2B and GAPM3, which link the IMC, alveolin network, and microtubules, are also dynamically phosphorylated.

Cluster 4, the only cluster characterized by decreasing phosphorylation, was functionally enriched in phosphoproteins involved in transporter and MAP kinase activity. ERK7 was dephosphorylated within 30 s of zaprinast treatment, whereas *Tg*TLK2 was dephosphorylated only at the final 60-s time point. ERK7 functions in parasite egress, motility, and invasion (O’Shaughnessy et al., 2020), whereas TLK2 regulates parasite replication (Smith et al., 2022; Varberg et al., 2018). Several phosphoproteins functioning in ion transport belonged to cluster 4, including a putative K^+^ voltage-gated channel complex subunit, a putative Vacuolar (H+)-ATPase G subunit, *Tg*ATP4, and the guanylyl cyclase (**Table 1**), which has a P-type ATPase domain with unknown ion specificity. Small-molecule transporters included a putative nucleoside transporter, ApiAT5-3, and a MFS family transporter. Cluster 4 was the only class of peptides that was not functionally enriched in Ca^2+^ binding proteins (**Figure 1E**).

We identified phosphoproteins with peptides belonging to several clusters (**Figure 1G**). Such proteins may have multiple phosphosites regulated with different kinetics by the same enzyme; or by different enzymes that alight upon the target at varying spatiotemporal scales. For example, SCE1 and TGGT1_309910 have phosphopeptides belonging to all three increasing clusters, likely resulting from phosphorylation by *Tg*CDPK3 and PKG, respectively. Indeed, SCE1 was implied to be a *Tg*CDPK3 target through a genetic suppressor screen (McCoy et al., 2017). TGGT1_309910 is the ortholog of *Plasmodium falciparum Pf*ICM1, a PKG substrate identified through proteomic interaction studies (Balestra et al., 2021), although it remains uncharacterized in *T. gondii*. By contrast, several proteins belong to both increasing and decreasing clusters, likely targets of both kinases and phosphatases. We consider these proteins candidate signaling platforms. Several such proteins localize to discrete domains of the apical complex, including the guanylyl cyclase; apical annuli proteins AAP2 and AAP5; apical polar ring proteins TGGT1_227000 and RNG2 (**Table 1**); the apical cap protein AC10 and a recently identified conoid gliding protein CGP. Along with the guanylyl cyclase, the cAMP-specific phosphodiesterase PDE2 and a Ca^2+^ influx channel with EF hands exhibit peptides belonging to both increasing and decreasing clusters. Our phosphoproteome thus identifies candidate mediators of the feedback between cyclic-nucleotide and Ca^2+^ signaling.

The waves of phosphoregulation largely corroborate the sequence of physiological events observed following zaprinast treatment. The first cluster likely includes the most proximal targets of PKG. Previous PKG-dependent phosphoproteomes from the related apicomplexan *P. falciparum* similarly implicate PI-PLC as a substrate of the kinase (Alam et al., 2015; Balestra et al., 2021; Brochet et al., 2014). The product of PI-PLC activity, IP_3_, stimulates Ca^2+^ store release in parasites (Garcia et al., 2017). Phosphoproteins in clusters 2, 3 and 4 likely contain the targets of Ca^2+^-regulated kinases and phosphatases.

### Thermal profiling identifies Ca^2+^-dependent shifts in *T. gondii* protein stability

The phosphoproteome identified numerous dynamic changes in response to Ca^2+^ release. However, this information alone is not sufficient to infer the enzymes responsible for phosphorylation, and which events may be functionally relevant. Thermal proteome profiling (TPP) has been used to identify small molecule–target interactions in living cells and cell extracts (Dai et al., 2019; Mateus et al., 2020; Savitski et al., 2014). TPP operates on the premise that ligand binding induces a thermal stability shift, stabilizing or destabilizing proteins that change conformationally in response to the ligand. Changes in stability can be quantified by MS (Dziekan et al., 2020; Reinhard et al., 2015). Cells are incubated with different concentrations of ligand and heated, causing thermal denaturation of proteins. The soluble protein is extracted and quantified with multiplexed, quantitative methods, giving rise to thousands of thermal denaturation profiles. Proteins engaging the ligand are identified by their concentration-dependent thermal shift. We previously used this method to identify the target of the antiparasitic compound ENH1 (Herneisen et al., 2020; Herneisen and Lourido, 2021) and to measure changes to the *T. gondii* proteome when depleting the mitochondrial protein DegP2 (Harding et al., 2020). In a conceptual leap, we reasoned that TPP could also detect Ca^2+^-responsive proteins if parasite extracts were exposed to different concentrations of Ca^2+^, allowing us to systematically identify the protein components of signaling pathways on the basis of biochemical interactions with Ca^2+^ and its effectors.

Intracellular free Ca^2+^ levels span three orders of magnitude, from low nanomolar in the cytoplasm to millimolar in organelles of the ER (Lourido and Moreno, 2015). To measure protein thermal stability at precisely defined [Ca^2+^]_free_, we combined crude parasite extracts with calibrated Ca^2+^ buffers in a solution mimicking the ionic composition of the cytoplasm. As a proof of principle, we measured the thermal stability of the Ca^2+^-dependent protein kinase *Tg*CDPK1, as the conformational changes of this enzyme have been structurally characterized (Ingram et al., 2015; Wernimont et al., 2010). Parasite lysates were adjusted to 10 different concentrations of free Ca^2+^ and heated to 58°C, based on prior estimates of the melting temperature of *Tg*CDPK1 (Herneisen et al., 2020). As measured by immunoblot band intensity, *Tg*CDPK1 was strongly stabilized by Ca^2+^ (**Figure 2A**), suggesting that our experimental system is sensitive to Ca^2+^-dependent stability changes. The calculated EC_50_ using this approach was in the low μM range, consistent with previous studies using recombinant enzymes (Ingram et al., 2015; Wernimont et al., 2010).

**Figure 2.**
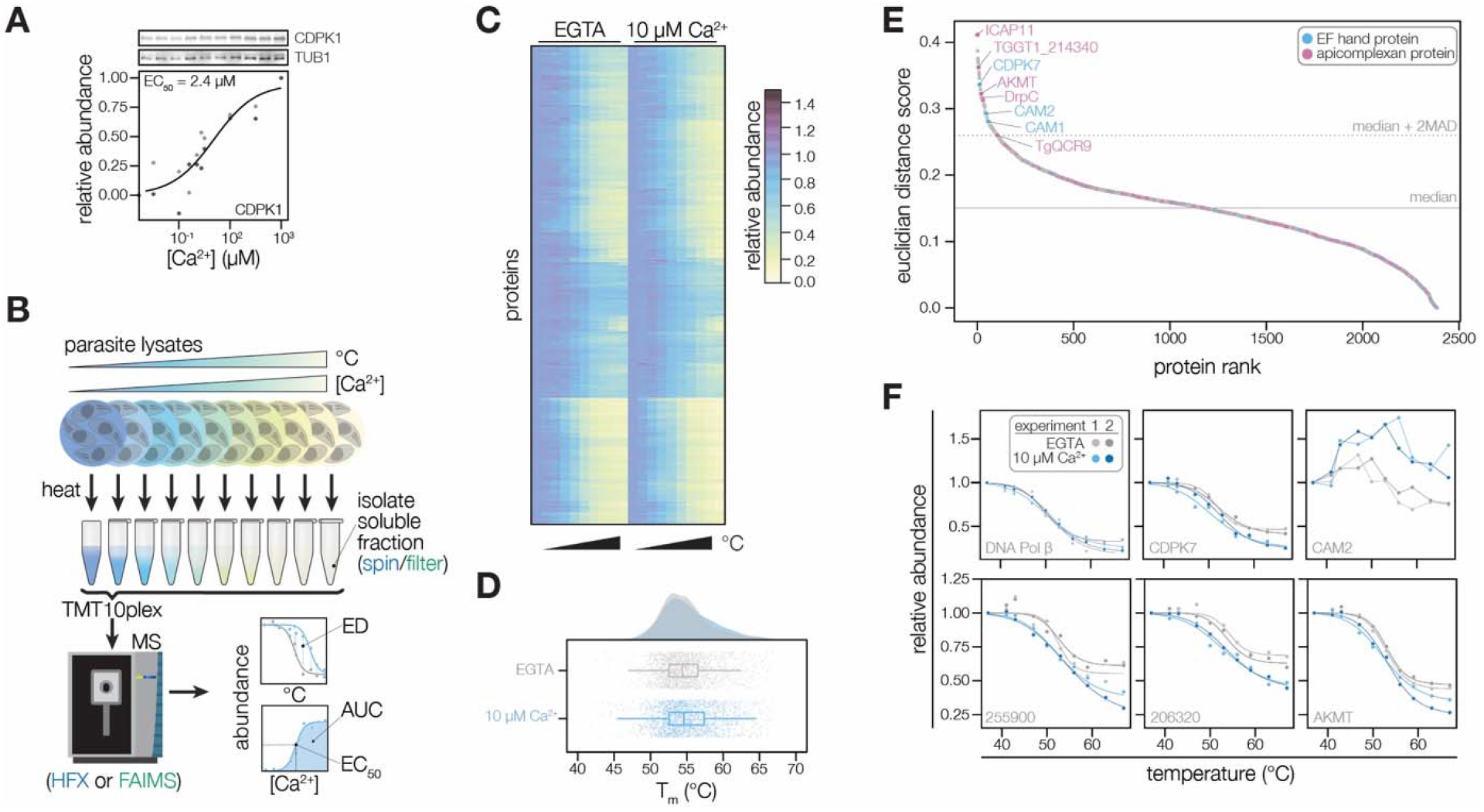
Thermal profiling identifies proteins that change stability in response to Ca^2+^. **(A)** Thermal shift assays can detect Ca^2+^-dependent stability of CDPK1 in extracts. Parasite lysates were combined with 10 concentrations of Ca^2+^ spanning the nanomolar to micromolar range. After denaturation at 58°C, the soluble fraction was separated by SDS-PAGE and probed for CDPK1. Band intensity was normalized to the no-Ca^2+^ control and was scaled. Points in shades of gray represent different replicates. A dose-response curve was calculated for the mean abundances. **(B)** Schematic of the thermal profiling workflow. In the temperature range experiment, parasite lysates were combined with EGTA or 10 μM [Ca^2+^]_free_ and heated at 10 temperatures spanning 37–67°C. In the concentration range experiment, parasite lysates were combined with 10 different [Ca^2+^]_free_ (nM-mM range) and heated at 50, 54, or 58°C. Temperature-range shifts were quantified by the euclidean distance (ED) score, a weighted ratio of thermal stability differences between treatments and replicates. Concentration-range shifts were summarized by pEC_50_, area under the curve (AUC), and goodness of fit (R^2^). **(C)** Heat map of protein thermal stability relative to the lowest temperature (37°C) in 0 or 10 μM Ca^2+^. The mean relative abundance at each temperature was calculated for 2,381 proteins. Proteins are plotted in the same order in both treatments. **(D)** Raincloud plots summarizing the distribution of T_m_ in lysates with EGTA (gray) or 10 μM [Ca^2+^]_free_ (blue). The average melting temperatures of proteins identified in two replicates were plotted. **(E)** Proteins rank-ordered by euclidean distance score quantifying the Ca^2+^-dependent shift in thermal stability. Solid and dotted lines represent the median ED score and two modified Z-scores above the median, respectively. Highlighted proteins have EF hand domains (blue) or are conserved in apicomplexans (pink). **(F)** Thermal profiles of individual proteins: DNA polymerase β (TGGT1_233820); the EF hand domain-containing proteins CDPK7 (TGGT1_228750) and the calmodulin-like protein CAM2 (TGGT1_262010); potential Ca^2+^-leak channels TGGT1_255900 and TGGT1_206320); and AKMT (TGGT1_216080).

The effect of Ca^2+^ on the global thermostability of the proteome has not been assessed. Therefore, we first generated thermal profiles of the *T. gondii* proteome without or with 10 μM Ca^2+^, which is representative of the resting and stimulated Ca^2+^ concentrations of cell cytoplasms (Lourido and Moreno, 2015). A thermal challenge between 37 and 67°C induced denatured aggregates, which were separated from stable proteins by ultracentrifugation. The soluble fraction was digested and labeled with isobaric mass tags, pooled, fractionated, and analyzed with an orbitrap mass spectrometer (**Figure 2B**). We detected 3,754 proteins, of which 2,381 yielded thermal profiles for both conditions in both replicates (**Figure 2C and Figure 2 Supplement**); the remaining proteins were not detected in all experiments. The median melting temperatures—the temperatures at which proteins are 50% denatured—were 54.4 and 54.7°C in the lysates without and with Ca^2+^, respectively. The distribution of melting temperatures was largely overlapping in the two conditions (**Figure 2D**), suggesting that Ca^2+^-dependent changes in protein stability were restricted to a subset of proteins. We additionally calculated an area under the curve (AUC) metric by numerical integration using the trapezoidal rule (Herneisen and Lourido, 2021) to compare the stabilities of proteins with atypical melting behavior (**Figure 2 Supplement**), such as components of the tubulin cytoskeleton or parasite conoid.

To discover proteins with Ca^2+^-dependent stability shifts in the initial temperature range experiment, we rank-ordered proteins by euclidean distance (ED) scores (Dziekan et al., 2020) quantifying the shift in thermal profiles with and without Ca^2+^ (**Figure 2B** and **2E**). The majority of proteins, such as DNA polymerase β (TGGT1_233820), exhibited similar melting behavior in both conditions (**Figure 2F**). Our analysis identified as Ca^2+^-responsive parasite-specific proteins with EF hands, including *Tg*CDPK7 and the calmodulin-like proteins CAM1 and CAM2 (**Table 1, Figure 2F** and **Figure 2 Supplement**). The ED metric identified the stability changes of both CAM proteins despite a lack of typical melting behavior, supporting the use of this statistic. Membrane proteins, including potential Ca^2+^-leak channels localizing to the ER (Barylyuk et al., 2020) (TGGT1_255900 and 206320), also exhibited thermal shifts (**Figure 2F**). Several proteins specific to the apicomplexan parasite phylum exhibited Ca^2+^-regulation (Barylyuk et al., 2020; Sidik et al., 2016a), including an apical lysine methyltransferase (**Figure 2F**) that relocalizes during Ca^2+^-stimulated egress (Heaslip et al., 2011); DrpC, which regulates the stability of parasite organelles; a hypothetical protein (TGGT1_214340) with no annotated domains (**Figure 2 Supplement**); and enzyme subunits involved in cellular metabolism.

The data also inform hypotheses about Ca^2+^ homeostasis and energetics in the parasite mitochondrion and apicoplast. A divergent subunit of the mitochondrial complex III, *Tg*QCR9, as well as components of the ATP synthase complex, were destabilized by Ca^2+^ (**Figure 2 Supplement**). These include the ATP synthase subunits alpha, gamma, 8/ASAP-15, and f/ICAP11/ASAP-10; and the ATP synthase-associated proteins ASAP-18/ATPTG14 and ATPTG1 (**Table 1**). TGGT1_209950, a conserved alveolate thioredoxin-like protein suggested to localize to the apicoplast by spatial proteomics (Barylyuk et al., 2020), was similarly destabilized by Ca^2+^ (**Figure 2 Supplement**). The apicomplexan mitochondrion has been reported to sense cytosolic Ca^2+^ fluctuations (Gazarini and Garcia, 2004), and the apicoplast uptakes Ca^2+^, perhaps through direct contact sites with the ER (Z.-H. Li et al., 2021). Furthermore, redox signals were recently reported to induce parasite Ca^2+^ signaling and motility (Alves et al., 2021; Stommel et al., 1997). Our resource identifies several proteins that may couple ion homeostasis and cellular metabolism. The recent structure of the apicomplexan ATP synthase suggests that all of the Ca^2+^-responsive elements identified here, including subunits restricted to apicomplexan parasites, protrude into the mitochondrial matrix (Mühleip et al., 2021), which has elevated Ca^2+^ (Giorgi et al., 2018). In the case of TGGT1_209950, plants and algae have Ca^2+^-sensing thioredoxins (calredoxins) in chloroplasts (Hochmal et al., 2016) with Ca^2+^-regulated activity mediated by EF hands (Charoenwattanasatien et al., 2020). TGGT1_209950 lacks EF-hands; however, structural modeling based on predicted secondary features (Meier and Söding, 2015) suggested similarity to calsequestrin, which lacks a structured Ca^2+^-binding motif. Our approach has thus identified novel candidate Ca^2+^-responsive proteins in metabolic organelles that would have been missed by bioinformatic searches.

### Determining the specificity and sensitivity of Ca^2+^-responsive proteins

The temperature range experiment has the advantage of generating complete thermal stability profiles, but does not inform the relative affinities of the Ca^2+^-dependent stability change. To address this gap, we examined protein stability across 10 Ca^2+^ concentrations (EGTA and 10 nM to 1 mM). Based on the temperature-range experiments, we selected thermal challenge temperatures of 50, 54, and 58°C to target protein with a range of thermal stabilities in these dose-response experiments (**Figure 2 Supplement**). We hypothesized that Ca^2+^-responsive proteins would exhibit sigmoidal dose-response trends in thermal stability, similarly to *Tg*CDPK1 (**Figure 2B**). We performed four independent concentration-range experiments on two mass spectrometers using different separation methods (ultracentrifugation or filtration) to capture different types of aggregates.

The concentration range thermal-profiling experiments provide insight into the magnitude of the Ca^2+^-dependent change (AUC parameter) and its dose-dependency (EC_50_) (**Figure 2B**). Clear responses were observed for several known Ca^2+^-binding enzymes with EF hands, including calmodulin, calcineurin, and several parasite Ca^2+^-dependent protein kinases (CDPKs; **Figure 3A**). These parameters can be used to classify proteins as stabilized (AUC > 1; e.g., *Tg*CDPK1, *Tg*CDPK2A, and *Tg*CDPK3) or destabilized (AUC < 1; e.g., calmodulin and calcineurin B) by Ca^2+^ (**Figure 3A**). Furthermore, EC_50_ measurements may inform specific hypotheses about a protein’s involvement in signaling or homeostasis, based on the Ca^2+^ concentration at which the protein is predicted to change.

**Figure 3.**
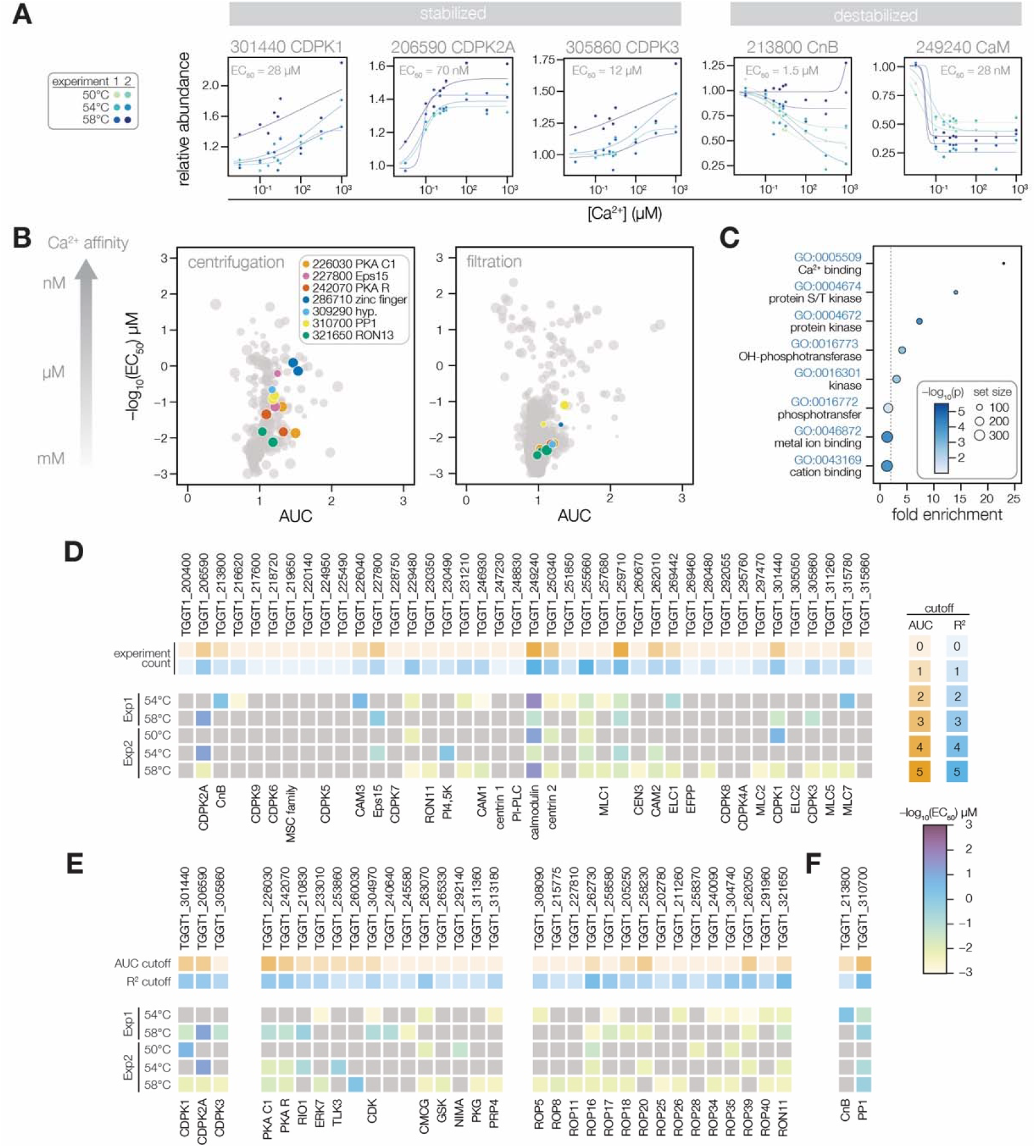
Thermal profiling identifies anticipated and unexplored Ca^2+^-responsive proteins. **(A)** Mass spectrometry-derived thermal profiles of EF hand-containing proteins stabilized or destabilized by Ca^2+^. Relative abundance is calculated relative to the protein abundance at 0 μM Ca^2+^. EC_50_ is the median of the EC_50_ values of the curves displayed on the plots. **(B)** The magnitude of Ca^2+^-dependent stabilization (AUC) plotted against the sensitivity (pEC_50_) for protein abundances exhibiting a dose-response trend with an R^2^ > 0.8. Point size is scaled to R^2^. Summary parameters for the different separation methods (ultracentrifugation or filtration) are plotted separately. Colors identify candidates with Ca^2+^-responsive behavior validated in Figure 4. **(C)** Gene ontology (GO) terms enriched among candidate Ca^2+^-responsive proteins (AUC greater than two modified Z scores and R^2^ dose-response > 0.8). Fold enrichment is the frequency of Ca^2+^-responsive proteins in the set relative to the frequency of the GO term in the population of detected proteins. Significance was determined with a hypergeometric test; only GO terms with *p* < 0.05 are shown. **(D–F)** EF hand domain proteins (D), protein kinases (E), and protein phosphatases (F) detected in the thermal profiling mass spectrometry datasets. The top rows indicate if a protein passed the AUC cutoff (orange) or R^2^ cutoff (blue) for dose-response behavior. The opacity of the band represents the number of experiments in which the protein exhibited the behavior (out of five). The five rows below summarize the pEC_50_ of each experiment in which the protein exhibited a dose-response trend with R^2^ > 0.8. Kinases are loosely grouped as CDPK’s (included as a reference), non-rhoptry kinases, and secretory pathway kinases.

Our results generated a *de novo* catalog of proteins with Ca^2+^-dependent stability. Of the 4,403 identified with sufficient quantification values for curve fitting, 228 were classified as Ca^2+^-responsive proteins based on exhibiting a dose-response trend (R^2^) greater than 0.8 with a stability change (AUC) of two modified Z-scores from the median (**Figure 3B, Figure Supplement 3**, and **Supplementary File 2**). Functional enrichment showed that such Ca^2+^-responsive proteins were significantly enriched in Ca^2+^-binding functions, protein kinase activity, and metal affinity (**Figure 3C**). We examined each of the 40 EF hand domain–containing proteins detected in our mass spectrometry experiments (**Figure 3D**). In total, 25 exhibited dose-responsive behavior. Signaling and trafficking proteins were tuned to respond at lower Ca^2+^ concentrations: *Tg*CDPK2A and CaM responded robustly with nM EC_50_; *Tg*CDPK1, *Tg*CDPK3, CnB, and Eps15 exhibited low μM EC_50_. In the cases of *Tg*CDPK1, CaM, and CnB, these values are consistent with the Ca^2+^ activation constants determined from studies with recombinant proteins (Feng and Stemmer, 2001, 1999; Wernimont et al., 2010). By contrast, myosin-associated subunits responded at higher levels: Myosin light chains (ELC1, MLC1, MLC5, and MLC7) and the calmodulin-like proteins CAM1 and CAM2 responded above 10 μM Ca^2+^ (**Table 1**). Two additional proteins (TGGT1_255660 and TGGT1_259710) consistently responded with μM EC_50_ values but have not been functionally characterized (**Figure 3 Supplement**).

Several signaling enzymes lacking canonical Ca^2+^-binding domains exhibited a dose-response trend (R^2^ > 0.8) in our thermal profiling experiments (**Figure 3B and 3E**). The PKA regulatory and catalytic subunits (TGGT1_242070 and TGGT1_226030) responded robustly to Ca^2+^ with 10–100 μM EC_50_ values in several experiments. A putative RIO1 kinase (TGGT1_210830) also responded consistently to Ca^2+^ concentration, although the magnitude of the change was small (**Figure 3 Supplement**). The atypical kinase ERK7 was destabilized at high Ca^2+^ concentrations (**Figure 3 Supplement**), similarly to other Ca^2+^-responsive proteins detected at the parasite apex. Rhoptry and dense granule kinases had EC_50_ values in between 100 μM and 1 mM, consistent with the high concentration of Ca^2+^ in the secretory pathway (**Figure 3E**). We also searched for Ca^2+^-responsive phosphatases. The known Ca^2+^-regulated phosphatase subunit, CnB, was destabilized (**Figure 3F**). Protein-phosphatase 1 (PP_1_) responded to Ca^2+^ more consistently than many known Ca^2+^-binding proteins (**Figures 3F** and **5A**), although the catalytic subunit is not an intrinsic Ca^2+^ sensor. Our resource thus places proteins without previously characterized Ca^2+^ responses within the broader Ca^2+^ signaling network.

Our search and annotation of Ca^2+^-responsive proteins also identified candidates that link Ca^2+^ and ion homeostasis. This list includes several transporters and channels (**Table 1**): *Tg*CAX, a protein with structural homology to LDL receptors (TGGT1_245610), two proteins with structural homology to the mitochondrial Ca^2+^ uniporter (TGGT1_257040 and TGGT1_211710), and an apicoplast two-pore channel. Several Ca^2+^-responsive hydrolases were also found, including SUB1, a metacaspase with a C2 domain, and a Ca^2+^-activated apyrase. These proteins may function during the intracellular, replicative phase of the lytic cycle, for which the function of Ca^2+^ is relatively unexplored.

Several divalent cations regulate protein structure and activity in addition to Ca^2+^, including Mg^2+^ and Zn^2+^. Such metal-binding proteins might appear Ca^2+^-responsive in our thermal profiling approach. We buffered our parasite extracts with excess Mg^2+^ (1 mM) to mitigate non-specific changes caused by the concentration of divalent cations. Nevertheless, to determine whether our approach identified additional metal-binding proteins, we compared the EC_50_ values of candidates that bind different divalent cations, as cataloged via the presence of Interpro domains and through manual annotation (**Figure 3 Supplement**). The EC_50_ values of putative Mg^2+^-binding proteins were mostly in the mM range, which may result from nonspecific displacement of Mg^2+^ by Ca^2+^. However, a subset of Mg^2+^-binding proteins were stabilized by micromolar Ca^2+^–for instance, glucosephosphate-mutase (GPM1) family proteins TGGT1_285980A and B. The GPM1 ortholog in *Paramecium*, parafusin (PFUS), is involved in Ca^2+^-regulated exocytosis (Subramanian and Satir, 1992), and the gene product in *T. gondii* has been observed to co-localize with micronemes in the apical third of the cell (Matthiesen et al., 2001). Therefore, the thermal stabilization of GPM1 could arise from Ca^2+^-dependent modifications to the enzyme (Subramanian et al., 1994; Subramanian and Satir, 1992).

### Validation of Ca^2+^-dependent thermal stability

We selected five proteins exhibiting consistent Ca^2+^-responsive behavior by MS for validation: an EF hand domain-containing protein (Eps15), two kinases (RON13 and PKA-C1), and two uncharacterized putative metal-binding proteins (TGGT1_286710 and TGGT1_309290). The first three candidates were selected for potential involvement in dynamic Ca^2+^-regulated processes. Eps15 (TGGT1_227800) was recently shown to mediate endocytosis in *P. falciparum* (Birnbaum et al., 2020) and localized to puncta bridging the inner membrane complex (IMC) and cytoskeleton in *T. gondii (Chern et al*., *2021)*. PKA-C1 is thought to antagonize Ca^2+^ signaling in *T. gondii (Jia et al*., *2017; Uboldi et al*., *2018b)*. RON13 is a secretory-pathway kinase that phosphorylates substrates in apicomplexan invasion-associated organelles called rhoptries (Lentini et al., 2021). The function of the remaining two proteins is unknown, although TGGT1_286710 contains a zinc finger domain and TGGT1_309290, annotated as a hypothetical protein, contains an HD/PDEase domain—both were indispensable for parasite survival (Sidik et al., 2016a).

We appended epitope tags to the endogenous locus of each candidate. These Ca^2+^-responsive proteins localized to diverse structures (**Figure 4B**). To validate the Ca^2+^-dependent stability of individual candidates, we prepared parasite extracts as described earlier, but relied on immunoblot readout instead of MS. In these five cases, stabilization of the candidates was confirmed in multiple biological replicates. Several controls (TUB1, MIC2, GAP45, and SAG1) could be shown to be stable across all Ca^2+^ concentrations tested (**Figure 4C** and **Figure 4 Supplement**). In the case of PKA-C1, Eps15, and both putative metal-binding proteins, the immunoblot experiments revealed an even higher EC_50_ than was measured in the MS experiments. We conclude that the stability changes detected by our global proteomics methods are robust, although the precise features of Ca^2+^-dependent stability may differ based on the method used to assess them.

**Figure 4.**
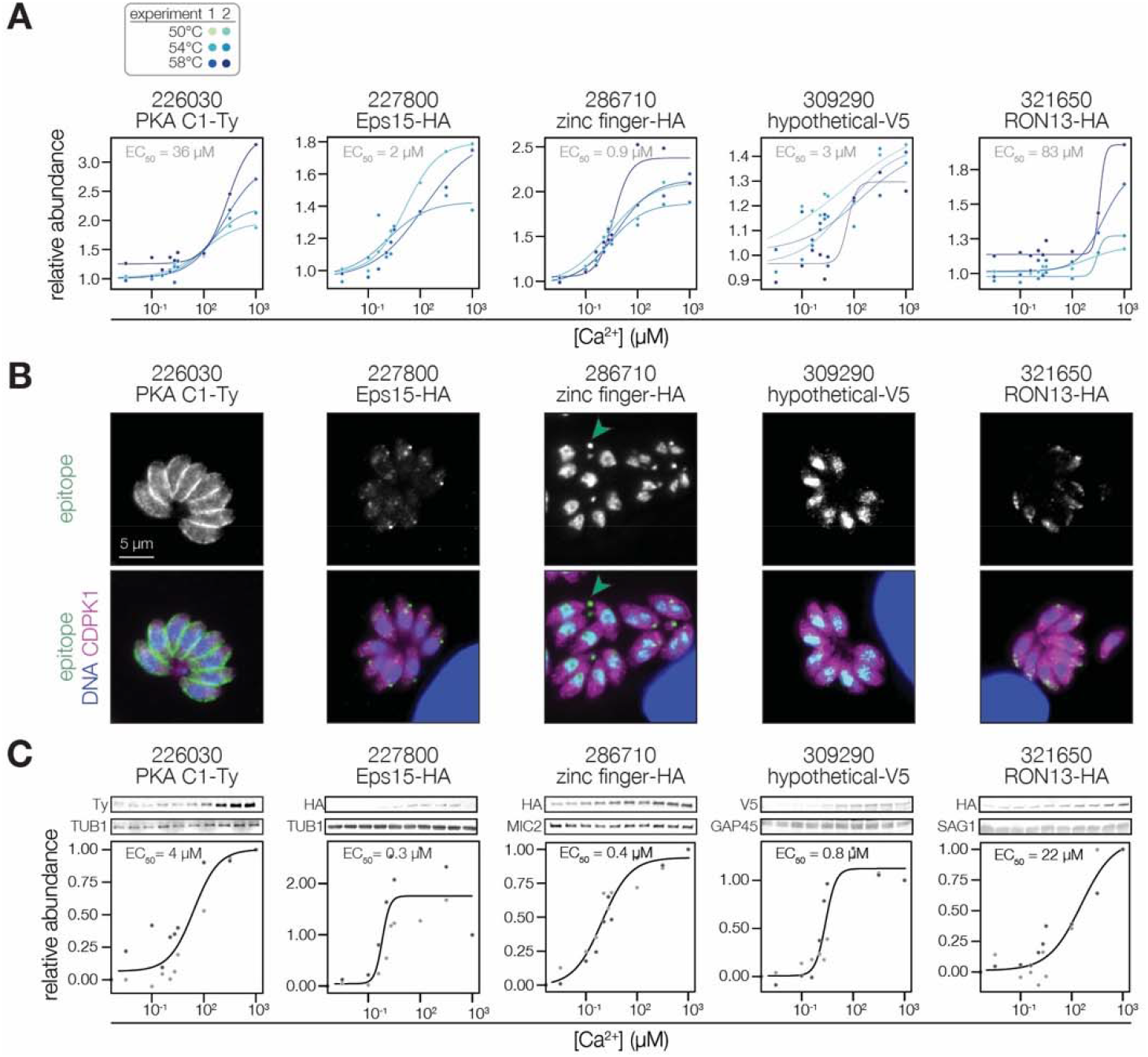
Validation of Ca^2+^-dependent thermal stability. **(A)** Mass spectrometry-derived thermal profiles of the candidates, as in Figure 3A. **(B)** Immunofluorescence images of fixed intracellular parasites expressing the indicated proteins with C-terminal epitopes at endogenous loci. Hoechst and anti-CDPK1 were used as counterstains in the merged image. Green arrowheads highlight an example of TGGT1_286710 basal pole staining. In the case of PKA C1/R, the stain of the R subunit is shown, as both subunits colocalize. **(C)** Immunoblot-derived thermal profiles of the candidates. Colors correspond to two independent replicates. Uncropped blots are shown in the **Figure 4 Supplement**.

### A PP_1_ holoenzyme serves Ca^2+^-responsive functions required for parasite spread

Our orthogonal proteomics approaches map Ca^2+^-responsive phosphorylation and thermal stability. Unexpectedly, the catalytic subunit of protein phosphatase 1 (PP_1_, TGGT1_310700) exhibited consistent stabilization by Ca^2+^ in our mass spectrometry experiments (**Figures 2B** and **5A**). The contribution of phosphatases to Ca^2+^ signaling in apicomplexans is poorly understood (Yang and Arrizabalaga, 2017) and the Ca^2+^-responsive behavior of PP_1_ has not been reported in other eukaryotes. The function of the phosphatase has not been directly studied in *T. gondii*, although PP_1_ inhibitors have been shown to block invasion of host cells (Delorme et al., 2002), and recent experiments in *P. falciparum* suggest that PP_1_ is required for the merozoite egress-to-invasion transition (Paul et al., 2020). Intriguingly, PP_1_ was recently observed to relocalize to the apical complex in highly motile *Plasmodium* berghei ookinetes (Zeeshan et al., 2021), suggesting that the enzyme may serve apicomplexan-specific, Ca^2+^-responsive functions in remodeling the parasite phosphoproteome.

To track localization during the *T. gondii* lytic cycle, we tagged the endogenous C terminus of PP_1_ with an mNG fluorophore and Ty epitope. We confirmed the Ca^2+^-dependent stability of PP_1_ by immunoblotting (**Figure 5B**). Live microscopy revealed a diverse array of PP_1_ localizations in parasites (**Figure 5C**). In accordance with imaging in *Plasmodium* (Zeeshan et al., 2021), PP_1_-mNG was distributed diffusely in the cytoplasm, as well as in foci resembling the nucleus, centrosome, and in some parasites, the periphery. These diverse localizations may arise from the association of PP_1_ with distinct regulatory subunits forming different functional holoenzymes, as characterized in metazoans (Brautigan and Shenolikar, 2018). To determine whether PP_1_ exhibits Ca^2+^-dependent relocalization, we treated parasites with zaprinast and A23187. The PP_1_ signal intensity increased at the apical cap and pellicle following zaprinast stimulation (**Figure 5C** and **Video 1**). However, Ca^2+^ ionophore treatment failed to induce the same relocalization patterns (**Figure 5C** and **Video 2**). The dynamics of PP_1_ activity thus appear specific to cGMP signaling within the parasites, which is nevertheless upstream of Ca^2+^ (Brown et al., 2016; Sidik et al., 2016b), and was similarly observed in *Plasmodium* (Paul et al. 2020).

**Figure 5.**
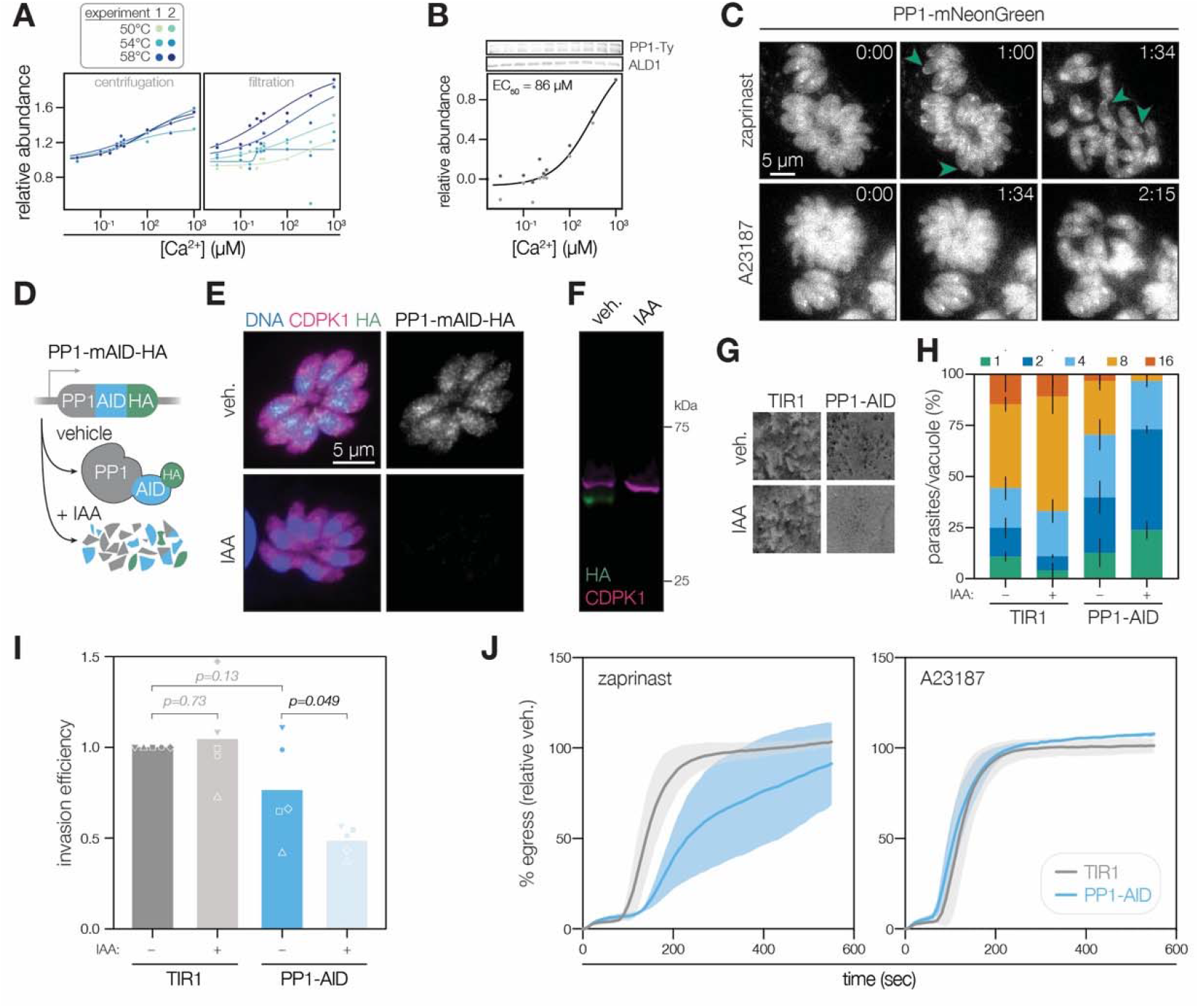
PP_1_ is a Ca^2+^-responsive enzyme involved in *T. gondii* egress and invasion. **(A)** The Ca^2+^-dependent stabilization of PP_1_ (TGGT1_310700) in each mass spectrometry experiment. **(B)** Immunoblotting for endogenously tagged PP_1_-mNG-Ty at different Ca^2+^ concentrations and thermal challenge at 58°C. Relative abundance is calculated relative to the band intensity at 0 μM Ca^2+^ and scaled. Points in shades of gray represent different replicates. A dose-response curve was calculated for the mean abundances. **(C)** Parasites expressing endogenously tagged PP_1_-mNG egress after treatment with 500 μM zaprinast or 4 μM A23187. Arrows show examples of PP_1_ enrichment at the apical end. Time after treatment is indicated in m:ss. **(D)** Rapid regulation of PP_1_ by endogenous tagging with mAID-HA. IAA, Indole-3-acetic acid (auxin). **(E)** PP_1_-mAID-HA visualized in fixed intracellular parasites by immunofluorescence after 3 hours of 500 μM IAA or vehicle treatment. Hoechst and anti-CDPK1 are used as counterstains (Waldman et al., 2020). **(F)** PP_1_-mAID-HA depletion, as described in **(E)**, monitored by immunoblotting. The expected molecular weights of PP_1_-mAID-HA and CDPK1 are 48 and 65 kDa, respectively. **(G)** Plaque assays of 1,000 TIR1 and PP_1_-mAID-HA parasites infected onto a host cell monolayer and allowed to undergo repeated cycles of invasion, replication, and lysis for 7 days in media with or without IAA. **(H)** The number of parasites per vacuole measured for PP_1_-mAID-HA and the TIR1 parental strain after 24 hours of 500 μM IAA treatment. Mean counts (*n* = 3) are expressed as a percentage of all vacuoles counted. **(I)** Invasion assays PP_1_-mAID-HA or TIR1 parental strains treated with IAA or vehicle for 3 hours. Parasites were incubated on host cells for 60 minutes prior to differential staining of intracellular and extracellular parasites. Parasite numbers were normalized to host cell nuclei for each field. Means graphed for *n* = 5 biological replicates (different shapes), Welch’s t-test. **(J)** Parasite egress stimulated with 500 μM zaprinast or 8 μM A23187 following 3 h of treatment with vehicle or IAA. Egress was monitored by the number of host cell nuclei stained with DAPI over time and was normalized to egress in the vehicle-treated strain. Mean ± S.D. graphed for *n* = 3 biological replicates.

The parasite apex and pellicle are hotspots for the signaling that potentiates motility and invasion, so the relocalization of PP_1_ suggests it may function in these processes. We created strains expressing the PP_1_ catalytic subunit with an endogenous C-terminal auxin-inducible degron (AID) for rapid conditional knockdown (**Figure 5D**). Conditional degradation of PP_1_ was confirmed by immunofluorescence and immunoblotting (**Figure 5E and 5F**). Parasites depleted of PP_1_ failed to form plaques (**Figure 5G**), implicating the phosphatase in one or more essential functions during the lytic cycle (Sidik et al., 2016a). Even in the absence of IAA, the PP_1_-AID strain formed small plaques, indicating substantial hypomorphism. PP_1_-AID parasites exhibited slower replication than untagged parasites, and this effect was exacerbated with IAA treatment (**Figure 5H**). Parasites depleted of PP_1_ had a reduced invasion efficiency (**Figure 5I**), although this effect was modest and subject to technical variation, likely arising from hypomorphism of the C-terminal tagged strain. Parasites treated with auxin egressed more slowly than untreated parasites when stimulated by zaprinast; however, egress kinetics were indistinguishable with a Ca^2+^ ionophore agonist (**Figure 5J**). Together, these results suggest that PP_1_ holoenzymes function at multiple steps in the lytic cycle. At least one holoenzyme relocalizes when parasite cGMP/Ca^2+^ pathways are stimulated. Because parasites lacking PP_1_ exhibit delays specifically in zaprinast-induced egress, we hypothesize that the peripheral holoenzyme enhances Ca^2+^ signaling; however, its requirement can be bypassed with nonspecific Ca^2+^ influx.

### A PP_1_ holoenzyme dephosphorylates signaling enzymes at the parasite periphery

PP_1_ is one of the major serine/threonine phosphatases in eukaryotic cells (Brautigan and Shenolikar, 2018). As the catalytic subunit relocalizes during cGMP/Ca^2+^-stimulated transitions in apicomplexans, we hypothesized that PP_1_ dephosphorylates crucial targets during egress and motility. To identify putative PP_1_ holoenzyme targets, we first treated intracellular PP_1_-AID parasites with IAA or vehicle for 3 hours prior to mechanically releasing parasites for analysis. We performed sub-minute phosphoproteomics by resuspending the extracellular parasites in a zaprinast solution followed by denaturation in SDS to stop enzymatic activity after 0, 10, 30, and 60 seconds. Multiplexing with TMTpro reagents followed by phosphopeptide enrichment allowed us to compare the zaprinast time courses with or without PP_1_-depletion in biological duplicate (**Figure 6A**). Analysis of an unenriched fraction of the proteome revealed significant depletion of PP_1_, which we confirmed in parallel by immunoblot (**Figure 6 Supplement**). The hypomorphism of the PP_1_-AID strain and reduced parasite yield resulted in a phosphoproteome with lower coverage: 6,916 phosphopeptides with quantification values. To identify peptides exhibiting PP_1_-dependent regulation, we selected peptides exhibiting a 2-fold difference in abundance between vehicle- and IAA-treated samples in both replicates for at least one time point. In total, 757 peptides passed this threshold.

**Figure 6.**
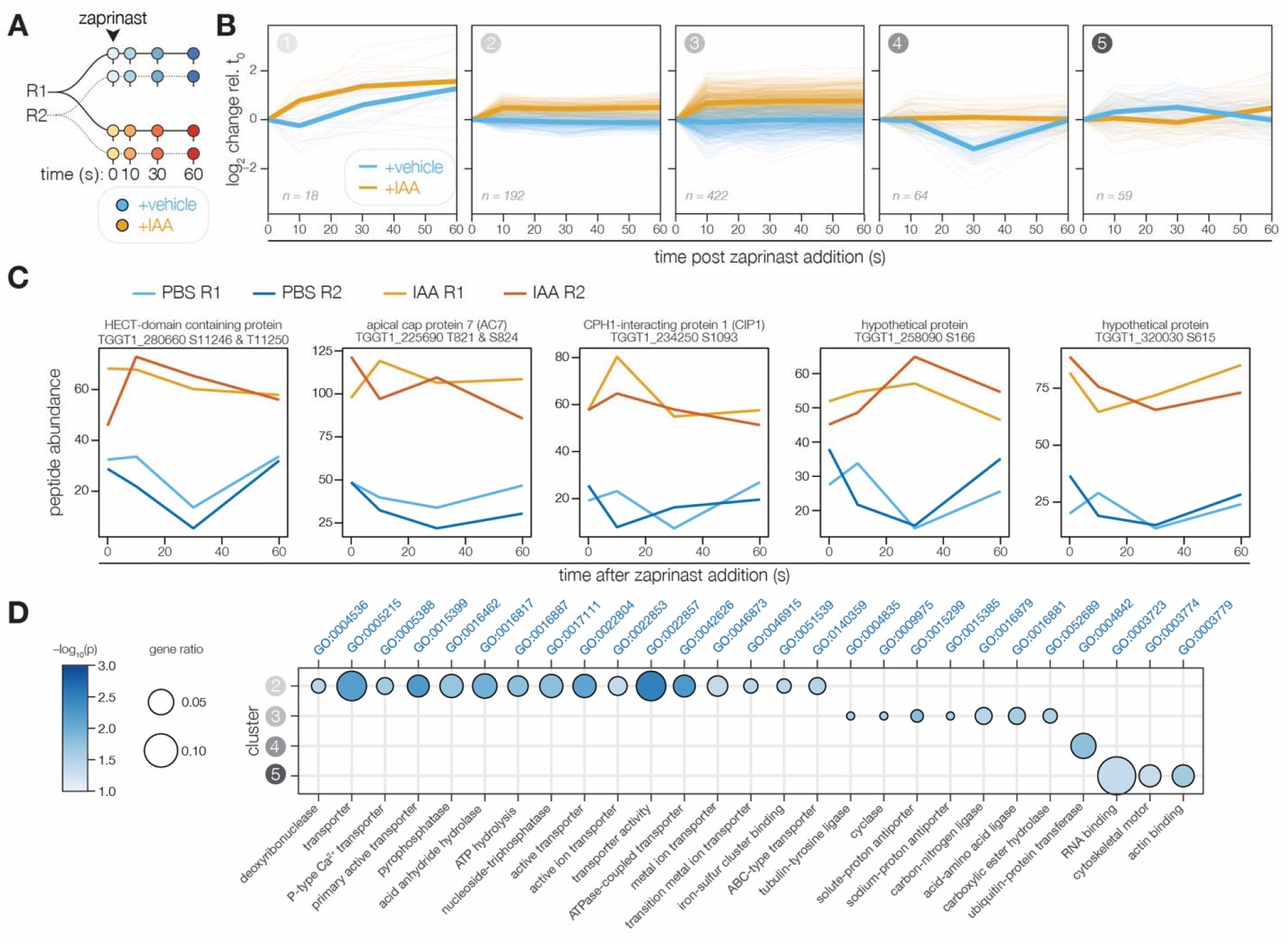
The PP_1_-dependent phosphoproteome. **(A)** Schematic of the phosphoproteomics time course. PP_1_-AID parasites were treated with IAA or vehicle for 3 hours. Extracellular parasites were then treated with zaprinast, and samples were collected during the first minute after stimulation. The experiment was performed in biological replicate (R1 and R2). **(B)** Five clusters were identified with respect to phosphopeptide dynamics and PP_1_-dependence. **(C)** Examples of phosphopeptides dynamically regulated by zaprinast and exhibiting PP_1_-dependent dephosphorylation. **(D)** GO terms enriched among phosphopeptides, grouped by cluster. Gene ratio is the proportion of proteins with the indicated GO term to the total number of proteins belonging to each cluster. Significance was determined with a hypergeometric test; only GO terms with *p* < 0.05 and represented by more than one protein are shown. Redundant GO terms were removed. Cluster 1 lacked enough peptides for enrichment analysis.

To focus on zaprinast-dependent changes, we clustered these peptides on the basis of their abundance relative to the earliest time point of stimulation. The peptides fit into five clusters with respect to PP_1_-dependence and kinetics (**Figure 6B**). Clusters 1, 2, and 3 contained peptides increasing in abundance upon stimulation in PP_1_-depleted parasites. The peptides belonging to Cluster 1 generally increase in abundance with zaprinast treatment, which occurs more rapidly with PP_1_ depletion. By contrast, the abundances of the 614 peptides belonging to clusters 2 and 3 were elevated in the absence of PP_1_, suggesting that PP_1_ antagonizes these phosphorylation events. Under normal conditions, peptides in cluster 4 decreased sharply in abundance between 10 and 30 seconds of zaprinast treatment and recovered by 60 seconds; however, these peptides did not change in abundance when PP_1_ was depleted. Peptides in cluster 5 increased gradually between 10 and 30 seconds and decreased to basal levels by 60 seconds; when PP_1_ is depleted, these peptides exhibit a delay.

The effect of PP_1_ disruption on the phosphoproteome was pervasive, which may reflect disruption of numerous PP_1_ holoenzymes. Many of the examined phosphopeptides exhibited substantial basal hyperphosphorylation in the absence of PP_1_ (**Figure 6C**). Given the likely pleiotropy of catalytic subunit depletion, we focused our analysis on perturbed pathways rather than individual targets (**Figure 6D**). Cluster 1 did not possess enough peptides for enrichment analysis. Cluster 2 was enriched in phosphoproteins functioning in transmembrane transport, including P-type ATPases and ABC transporters. Cluster 3 contained both the guanylyl and adenylyl cyclases and was further enriched in putative sodium-hydrogen exchangers and tubulin-tyrosine ligases. Cluster 4 was solely enriched for proteins involved in ubiquitin transfer, including TGGT1_280660, an uncharacterized HECT domain-containing protein (**Figure 6C**). Cluster 5 phosphoproteins were notably involved in cytoskeletal motor activity, actin binding, and RNA binding. Numerous apical proteins exhibited PP_1_-dependent phosphorylation (**Table 1**), including AC7, CIP1, and two hypothetical proteins—TGGT1_258090 and TGGT1_320030, localizing to the conoid base and the second apical polar ring, respectively (Koreny et al., 2021). The protein abundances did not vary between vehicle and IAA treatment, indicating that the phosphopeptide abundance changes arose from dynamic covalent modifications. The PP_1_-dependent phosphoproteome therefore supports the existence of apical and peripheral PP_1_ holoenzymes, as observed by live microscopy.

### PP_1_ activity is important for Ca^2+^ entry to enhance the kinetics of egress

The phosphoproteomics data pointed to ion homeostasis dysregulation in the absence of PP_1_ (**Figure 6D**). Numerous transporters are hyperphosphorylated when PP_1_ is depleted, including Ca^2+^ ATPases, proton transporters, and MFS and ABC-family transporters. We hypothesized that aberrant Ca^2+^ mobilization may underlie the phenotypes observed in the motile stages of the lytic cycle upon PP_1_ depletion (**Figure 5**), as Ca^2+^ release and signaling are required for egress and invasion. To monitor Ca^2+^ mobilization in parasites, we introduced the genetically encoded Ca^2+^ indicator GCaMP6f into a defined chromosomal locus (Chen et al., 2013; Herneisen et al., 2020). By tracking the vacuole fluorescence of PP_1_-depleted parasites, we observed delayed Ca^2+^ mobilization and egress following zaprinast treatment (**Figure 7A–C, Figure 7 Supplement**, and **Video 1**). Despite the delay, IAA-treated parasites attained the same increase in Ca^2+^ levels prior to egress as vehicle-treated parasites (**Figure 7 Supplement**). By contrast, Ca^2+^ mobilization upon A23187 treatment was unperturbed in PP_1_-depleted parasites (**Figure 7D–F, Figure 7 Supplement** and **Video 2**). Motivated by recent experiments suggesting that *T. gondii* exhibits Ca^2+^-activated Ca^2+^ entry upon zaprinast treatment (Vella et al., 2021), we loaded PP_1_-AID parasites with the ratiometric Ca^2+^ indicator Fura2-AM and observed reduced Ca^2+^ entry (**Figure 7G and 7H**). The integration of our global datasets therefore identified PP_1_ as a Ca^2+^-responsive enzyme and precipitated discovery of its role in Ca^2+^ entry.

**Figure 7.**
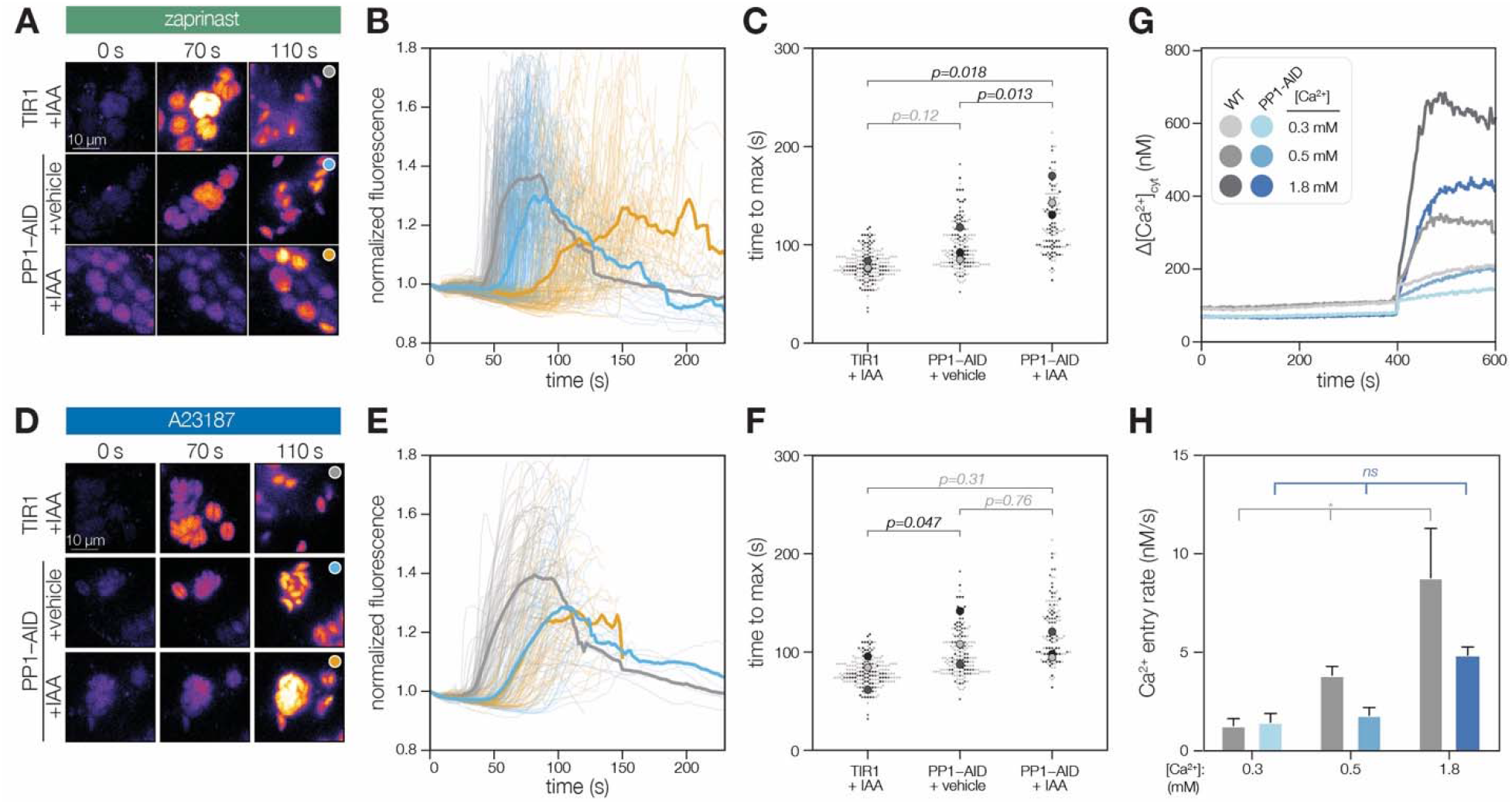
PP_1_ function ensures rapid Ca^2+^ mobilization prior to zaprinast-induced egress. **(A)** Selected frames of time-lapse images of PP_1_-AID parasites expressing the genetically encoded Ca^2+^ indicator GCaMP following treatment with 500 μM zaprinast. **(B)** Normalized GCaMP fluorescence of individual vacuoles was tracked after zaprinast treatment and prior to egress (transparent lines). The solid line represents the mean normalized fluorescence of all vacuoles across *n = 3* biological replicates. **(C)** The time to maximum normalized fluorescence of individual vacuoles after zaprinast treatment. Different replicates are shown in different shades of gray. Transparent dots represent individual vacuoles; solid dots are the mean for each replicate. P-values were calculated from a two-tailed t-test. **(D)** Selected frames of time-lapse images of PP_1_-AID parasites expressing the genetically encoded Ca^2+^ indicator GCaMP following treatment with 4 μM A23187. **(E)** Normalized GCaMP fluorescence of individual vacuoles was tracked after A23187 treatment and prior to egress (transparent lines). The solid line represents the mean normalized fluorescence of all vacuoles across *n = 3* biological replicates. **(F)** The time to maximum normalized fluorescence of individual vacuoles after A23187 treatment. Different replicates are shown in different shades of gray. Transparent dots represent individual vacuoles; solid dots are the mean for each replicate. P-values were calculated from a two-tailed t-test. **(G)** Addition of extracellular Ca^2+^ at different concentrations to Fura2/AM-loaded parasites leads to intracellular Ca^2+^ increase in TIR1 parasites treated with IAA for 5 hours. The response of the PP_1_-AID parasites treated with IAA was significantly reduced. The buffer contained 100 mM EGTA and Ca^2+^ was added at the indicated concentration at the indicated time. Representative experiment from at least three biological replicates. **(H)** The slope of the trace at the time of addition of Ca^2+^ was measured as the change in the concentration of Ca^2+^ during the initial 20 s after addition of Ca^2+^. Each bar represents the average of a minimum of three biological replicates. ANOVA was used for the statistical analyses. * p< 0.01.

## DISCUSSION

Apicomplexan signaling pathways have largely been characterized through a combination of genetic manipulation, pharmacological perturbation, and physiological observation (Bisio and Soldati-favre, 2019; Brown et al., 2020; Lourido and Moreno, 2015). Such approaches have been sufficient to document feedback loops between cyclic nucleotide signaling and Ca^2+^-store release and influx. However, the molecular mechanisms linking these pathways have remained obscure, as genomic annotation or bioinformatic analysis are complicated by the evolutionary divergence of apicomplexans. Unbiased, functional proteomics approaches have allowed us to map a Ca^2+^-responsive proteome de novo. Our resource places proteins already under study within Ca^2+^ signaling pathways and identifies hubs serving as pressure points for the perturbation of multiple signaling processes.

Our time-resolved phosphoproteomic data adds molecular depth to known changes in parasite physiology. The first cluster of phosphoproteins to respond to zaprinast stimulation includes second messenger amplifiers: cyclic nucleotide phosphodiesterases (PDE1 and PDE2); lipid kinases, phosphatases, and phospholipases; and the guanylyl cyclase (Bisio et al., 2019; Brown and Sibley, 2018). This cluster also contains early Ca^2+^ sensors and transducers, such as *Tg*CDPK2A and *Tg*CDPK7 (Bansal et al., 2021), which additionally responded to Ca^2+^ in our thermal profiling experiments. The next two clusters reveal rewiring of small molecule homeostasis, with non-Ca^2+^ channels and transporters phosphorylated first. Parasites have long been reported to respond to extracellular H^+^ and K^+^ levels (Moudy et al., 2001; Roiko and Carruthers, 2013), although the proteins responsible, and sequence of events transducing signals across the parasite plasma membrane, remained obscure. Our proteomic datasets identified dynamic phosphorylation of ATPase2, NHE1, and two uncharacterized putative Ca^2+^-activated K^+^ channels. ATPase2 was recently reported to mediate essential lipid homeostasis at the parasite plasma membrane (Chen et al., 2021). Mutants lacking NHE1 exhibited elevated cytosolic Ca^2+^, yet were less susceptible to Ca^2+^ ionophore-induced egress (Arrizabalaga et al., 2004), suggesting that monovalent cation and Ca^2+^ homeostasis mechanisms are intertwined. Indeed, the third wave of phosphorylation targets Ca^2+^ and other divalent cation ATPases, including *Tg*A1, *Tg*CuTP, and other P-type ATPases with uncharacterized ion specificity. Transporters belonging to these clusters are noted *Tg*CDPK3 substrates, including ATP4, SCE1, and ApiAT5-3; disruption of these transporters failed to alter cytosolic Ca^2+^ levels. A cluster of dynamic dephosphorylation events highlights the need for phosphatases to reverse modifications as parasites transition from one phase to another (Brautigan and Shenolikar, 2018; Yang and Arrizabalaga, 2017). These phosphoproteins were also enriched for channel or transporter functions, and in several cases also contained sites regulated by kinases. Apicomplexans must maintain homeostasis as ion gradients are inverted when parasites transition from intracellular to extracellular, likely requiring extreme rewiring of cellular physiology. Our phosphoproteomics dataset reveals that parasite ion homeostasis pathways are hard-wired into second messenger signal transduction, perhaps in anticipation of environmental challenge.

We can infer mechanisms of crosstalk between second messenger signaling networks, different classes of enzymes, and post-translational modifications from our global proteomic profiles. Proteins regulated by several kinases and phosphatases with different kinetics, may function to concentrate these enzymes in local signaling hotspots acting as signaling scaffolds. Several of these dynamically regulated proteins themselves contain second messenger–sensing domains, such as EF hands (TGGT1_216620 and *Tg*CDPK2A) and cyclic nucleotide–binding domains (PDE2 and AAP2). Intriguingly, both subunits of PKA, the only known cAMP receptor in *T. gondii*, are stabilized by Ca^2+^. PKA has been proposed to negatively regulate Ca^2+^ signaling in parasites (Jia et al., 2017; Uboldi et al., 2018a). Here, we additionally observe Ca^2+^ regulating PKA, although the molecular basis of this stability change remains to be established. Finally, the stimulated phosphoproteome exhibits extensive crosstalk with GTPase regulation and ubiquitin transfer pathways, suggesting that there remain vast and unexplored signaling networks, in addition to phosphorylation, that remodel cellular states in *T. gondii*.

The Ca^2+^ responsiveness of PP_1_ revealed by our TPP analysis was surprising. In most organisms, the dominant Ca^2+^-regulated phosphatase is calcineurin (Shi, 2009). Although PP_1_ holoenzymes have been reported to modulate Ca^2+^ channels, the Ca^2+^-responsive behavior of PP_1_ has not been reported in any other eukaryote. The function of the phosphatase had not been directly studied in *T. gondii*, although PP_1_ inhibitors have been shown to block invasion of host cells (Delorme et al., 2002) and recent experiments in *P. falciparum* suggest that PP_1_ is required for an as-yet undetermined step in the kinetic phase of infection (Paul et al., 2020). Guided by our global experiments, we endogenously tagged the C-terminus of PP_1_ and observed diverse localization patterns.

Numerous regulatory subunits control the activity, localization, and specificity of the single catalytic subunit of the phosphatase (Yang and Arrizabalaga, 2017), giving rise to functionally distinct PP_1_ holoenzymes that play conserved roles in cell cycle progression (Bertolotti, 2018). Stimulating the transition from the replicative to the kinetic phase by treating parasites with the phosphodiesterase inhibitor zaprinast led to rapid enrichment of PP_1_ at the parasite periphery and apical end, hotspots of parasite signaling. Notably, PP_1_ was recently reported to relocalize to the apical complex in *Plasmodium* berghei ookinetes (Zeeshan et al., 2021). Although the proposed role of PP_1_ in ookinetes was to set cell polarity, given our phenotypic observations in *Toxoplasma*, we propose instead that the apical PP_1_ holoenzyme serves apicomplexan-specific, Ca^2+^-responsive functions supporting the parasitic lifestyle (Park et al., 2019; Paul et al., 2015; Philip and Waters, 2015). Indeed, parasites conditionally depleted of PP_1_ exhibited reduced invasion and delayed host cell lysis in response to zaprinast.

The parasite PP_1_ holoenzyme is as yet undefined. A recent phosphoproteome of *P. falciparum* parasites depleted of PP_1_ identified ∼5,000 phosphosites from ∼1,000 proteins (Paul et al., 2020). Despite complications in determining direct substrates, the study proposed PP_1_ regulation of a HECT E3 ligase and guanylate cyclase upstream of PKG, providing a potential functional link between PP_1_ and motility. Our experimental design benefited from higher temporal resolution and the ability to directly trigger the signaling pathways in question, providing a more comprehensive analysis of phosphatase activity during stimulated motility. Our analysis uncovered two patterns of PP_1_-dependent regulation: (i) sites hyperphosphorylated only in the absence of PP_1_ (clusters 2 and 3), and (ii) zaprinast-dependent sites no longer regulated in the absence of PP_1_ (cluster 4). The guanylyl cyclase and possible ortholog of the HECT E3 ligase identified in the *P. falciparum* study both included PP_1_-dependent phosphosites in our experiment, suggesting that these PP_1_-regulated cascades are important and conserved in apicomplexan parasites. The guanylyl cyclase belongs to one of the hyperphosphorylated clusters. If guanylyl cyclase function is compromised in the absence of PP_1_, accumulation of cGMP to the levels required for egress may be delayed following inhibition of phosphodiesterases, partially explaining the delayed zaprinast response observed in PP_1_-depleted parasites (Paul et al., 2020). The set of phosphoproteins normally dephosphorylated by PP_1_ during zaprinast treatment was uniquely enriched in ubiquitin-transfer enzymes. Prior proteomic enrichment studies observed an abundance of ubiquitin modifications in cell cycle-regulated proteins, including the peripheral parasite inner membrane complex (Silmon de Monerri et al., 2015). However, the ubiquitination pathway is largely unexplored in the context of apicomplexan motility. HECT E3 ligase inhibitors reduced merozoite egress in *P. falciparum* (Paul et al., 2020). Our proteomic datasets reveal that the phosphorylation states of ubiquitin transferases and hydrolases change within 60 seconds of cGMP elevation, suggesting that dynamic ubiquitination intersects with dynamic phosphorylation to regulate transitions between intracellular and extracellular states.

The enrichment of transporters in the basally dysregulated sites motivated us to explore PP_1_ phenotypes associated with Ca^2+^ mobilization and entry. We observed that PP_1_ activity enhances Ca^2+^ entry, which itself enhances parasite motility (Pace et al., 2014; Vella et al., 2021). Synthesizing the phosphoproteomic, phenotypic, and thermal profiling data, which reports a micromolar-range EC_50_ of PP_1_ for Ca^2+^, we propose that the Ca^2+^-regulated PP_1_ holoenzyme assembles following an initial wave of Ca^2+^ store release, and then combats negative feedback mechanisms promoting Ca^2+^ uptake. Such negative feedback loops between Ca^2+^-stimulated kinases and Ca^2+^ uptake have been reported for *Tg*CDPK1 and *Tg*CDPK3 (Herneisen et al., 2020; Stewart et al., 2017). These results argue that, like calcineurin, PP_1_ plays an unequivocal role in the Ca^2+^ signaling network of *T. gondii*, and together both phosphatases are likely responsible for much of the Ca^2+^-regulated serine/threonine phosphatase activity in parasites.

The Ca^2+^ signaling field has benefited from decades of research in other model organisms, in which pathway connections were incrementally extended based on prior knowledge about the system (Berridge et al., 2000; Clapham, 2007; Luan and Wang, 2021). In contrast to this paradigm, we developed a global mass spectrometry-based approach to systematically identify the protein components of Ca^2+^ signaling pathways on the basis of dynamic phosphoregulation and biochemical thermal stability. We applied this technology to apicomplexans, which are widespread eukaryotic human parasites whose signaling pathways remain largely unmapped due to their evolutionary divergence from model organisms (Lourido and Moreno, 2015). In principle, this approach can be applied to other post-translational modifications, natural ligands (Lim et al., 2018; Sridharan et al., 2019), and organisms (Jarzab et al., 2020) to establish—and in some cases re-evaluate—the topology of complex signaling pathways.

## METHODS

### Reagents and cell culture

*T. gondii* parasites of the type I RH strain Δ*ku80*/Δ*hxgprt* genetic background (ATCC PRA-319) were grown in human foreskin fibroblasts (HFFs, ATCC SCRC-1041) maintained in DMEM (GIBCO) supplemented with 3% inactivated fetal calf serum (IFS) and 10 μg/mL gentamicin (Thermo Fisher Scientific). When noted, DMEM was supplemented with 10% IFS and 10 μg/mL gentamicin.

### Parasite transfection and strain construction

#### Genetic background of parasite strains

Existing *T. gondii* RH strains were used as genetic backgrounds for this study. All strains contain the *Δku80Δhxgprt* mutations to facilitate homologous recombination (Huynh and Carruthers, 2009). TIR1/RH expresses the TIR1 ubiquitin ligase (Brown et al., 2017). TIR1/pTUB1-GCaMP additionally expresses a GCaMP6f transgene (Smith et al., 2022). DiCre_T2A/RH uses the same promoter to express both subunits of Cre, separated by T2A skip peptides (Hunt et al., 2019).

#### TIR1/pMIC2-MIC2Gluc-P2A-GCaMP6f

A new TIR1/RH strain expressing GCaMP6f under different regulatory elements was constructed as a part of this study. The sequence pMIC2_MIC2Gluc_myc_P2A_GCaMP6f_DHFR3 UTR was amplified with primers P1 and P2 from plasmid Genbank ON312866 to yield a repair template with homology to the 5 and 3 ends of a defined, neutral genomic locus (Markus et al., 2019). Approximately 1 × 10^7^ extracellular TIR1 parasites were transfected with 20 μg gRNA/Cas9-expression plasmid pBM041 (GenBank MN019116.1) and 10 μg repair template. GFP-positive clones were isolated by limiting dilution following fluorescence-activated cell sorting.

#### Endogenous tagging of candidate genes

Genes were endogenously tagged using the previously described high-throughput (HiT) tagging strategy (Smith et al., 2022). Cutting units specific to each candidate were purchased as IDT gBlocks (P3-P7) and assembled with the following empty HiT vector backbones: pALH083 (V5-mCherry-mAID-Ty; GenBank: ON312867), pALH052 (V5-mCherry-mAID-HA; GenBank: OM863784), pALH047 (V5-3HA; GenBank: ON312868), pGL015 (V5-mNG-mAID-Ty; GenBank OM640005), pEC341 (HA-loxP-U1; GenBank: OM863788), and pALH086 (V5-mAID-HA; GenBank ON312869).

Between 20 and 50 μg of each vector was BsaI-linearized and co-transfected with 50 μg of the pSS014 Cas9-expression plasmid (GenBank: OM640002). Vectors targeting TGGT1_226030 (PKA-C1) and TGGT1_227800 (Eps15) were transfected into TIR1/pTUB1-GCaMP or TIR1/pMIC2-MIC2Gluc-P2A-GCaMP6f. Vectors targeting TGGT1_286710 and TGGT1_309290 were transfected into TIR1/RH. Vectors targeting TGGT1_321650 were transfected into DiCre/RH. Parasite populations were selected for 1 week in standard media with 1 μM pyrimethamine or 25 μg/mL mycophenolic acid and 50 μg/mL xanthine, followed by subcloning into 96-well plates. Single clones were screened for tag expression by immunofluorescence and immunoblot.

#### Endogenous tagging of PP_1_

A cutting unit specific to the C-terminus of PP_1_ (TGGT1_310700; P8) was assembled into the pALH086 HiT vector backbone. Approximately 50 μg of this vector was BsaI-linearized and co-transfected with 50 μg of the pSS014 Cas9-expression plasmid into TIR1/RH or Parasite populations were selected for 1 week in standard media with 25 μg/mL mycophenolic acid and 50 μg/mL xanthine and were then subcloned by limiting dilution. Single clones were screened for tag expression by immunofluorescence, immunoblot and sequencing of the junction spanning the 3 CDS and the tag.

### Sub-minute phosphoproteomics experiments

#### Parasite harvest and treatment

*T. gondii* tachyzoites from the RH strain were infected onto confluent HFF monolayers in 4–8 15-cm dishes. After the parasites had completely lysed the host cell monolayer (40-48 hours post-infection), extracellular parasites were passed through 5 μm filters into 50 ml conical vials. The samples were spun at 1000 x g for 7 minutes. The supernatant was decanted, and the parasite pellet was resuspended in 1 ml FluoroBrite DMEM lacking serum and transferred to a 1.5 ml protein low-bind tube. The sample was spun in a mini centrifuge at 1000 x g for 7 minutes. The supernatant was aspirated, and the parasite pellet was resuspended in 800 μl FluoroBrite DMEM. The sample was split into aliquots of 300 and 500 μl, which were spun at 1000 x g for 7 minutes followed by aspiration of the supernatant. The pellet containing ⅝of the parasite harvest was resuspended in 250 μl 500 μM zaprinast in FluoroBrite DMEM. Aliquots of 50 μl were removed and combined with 50 μl 2X lysis buffer (10% SDS, 4 mM MgCl_2_, 100 mM TEAB pH 7.55 with 2X Halt Protease and Phosphatase Inhibitors and 500 U/ml benzonase) at 0, 5, 10, 30, and 60 seconds post-stimulation. The final composition of the lysate was 5% SDS, 2 mM MgCl_2_, 50 mM TEAB pH 8.5 with 1X Halt Protease and Phosphatase Inhibitors and 250 U/ml benzonase. The pellet containing ⅜of the parasite harvest was resuspended in 150 μl 0.5% DMSO in FluoroBrite DMEM. Aliquots of 50 μl were removed and combined with 50 μl 2X lysis buffer at 0, 10, and 30 seconds post-stimulation.

#### Protein cleanup and digestion

Proteins were cleaned up and digested using a modified version of the S-trap protocol (Protifi). Proteins in the lysates were reduced with the addition of 5 mM TCEP and heating at 55°C for 20 minutes. Alkylation was performed for 10 minutes at room temperature with 20 mM MMTS. The lysates were acidified to a final concentration of 2.5% v/v phosphoric acid. A 6X volume of S-trap binding buffer (90% methanol, 100 mM TEAB pH 7.55) was added to each sample to precipitate proteins. The solution was loaded onto S-trap mini columns (Protifi) and spun at 4,000 x g until all of the solution had been passed over the column. The columns were washed four times with 150 μl S-trap binding buffer, followed by a 30 second spin at 4,000 x g. Proteins were digested overnight at 37°C in a humidified incubator with 20 μl of 50 mM TEAB pH 8.5 containing 2 μg of trypsin/LysC mix (Thermo Fisher Scientific). Peptides were eluted in three 40 μl washes with 50 mM TEAB, 0.2% formic acid, and 50% acetonitrile/0.2% formic acid. The eluted peptides snap-frozen and lyophilized.

#### TMTpro labeling

The dried peptides were resuspended in 50 μl 100 mM TEAB 8.5. The peptide concentrations of 1/50 dilutions of the samples were quantified using the Pierce Fluorometric Peptide Assay according to manufacturer’s instructions. Sample abundances were normalized to 50 μg peptides in 50 μl 100 mM TEAB pH 8.5. Each sample was combined with TMTpro reagents at a 5:1 label:peptide weight/weight ratio. Labeling reactions proceeded for 1 hour at room temperature with shaking at 350 rpm. Unreacted TMT reagent was quenched with 0.2% hydroxylamine. The samples were pooled, acidified to 3% with formic acid, and were loaded onto an EasyPep Maxi Sample Prep column. The samples were washed and eluted according to the manufacturer’s instructions. 5% by volume of the elution was reserved as an unenriched proteome sample. The eluted peptides were snap-frozen and lyophilized until dry.

#### Phosphopeptide enrichment

Phosphopeptides were enriched using the SMOAC protocol (Tsai et al., 2014). Resuspended TMTpro-labeled samples were enriched with the High-Select™ TiO2 Phosphopeptide Enrichment Kit (Thermo Fisher Scientific A32993) according to the manufacturer’s instructions. The flow-through and the eluate containing phosphopeptides were immediately snap-frozen and lyophilized. The flow-through was resuspended and enriched with the High-Select™ Fe-NTA Phosphopeptide Enrichment Kit (Thermo Fisher Scientific A32992) according to the manufacturer’s instructions. The eluted phosphopeptides were immediately snap-frozen and lyophilized.

#### Fractionation

Unenriched and enriched proteome samples were fractionated with the Pierce High pH Reversed-Phase Peptide Fractionation Kit according to the manufacturer’s instructions for TMT-labeled peptides. The phosphopeptides enriched with the TiO2 and Fe-NTA kits were combined prior to fractionation.

#### MS data acquisition

The fractions were lyophilized and resuspended in 10-20 μl of 0.1% formic acid for MS analysis and were analyzed on an Exploris 480 Orbitrap mass spectrometer equipped with a FAIMS Pro source (Bekker-Jensen et al., 2020) connected to an EASY-nLC chromatography system as described above. Peptides were separated at 300 nl/min on a gradient of 5–20% B for 110 minutes, 20–28% B for 10 minutes, 28–95% B for 10 minutes, 95% B for 10 minutes, and a seesaw gradient of 95–2% B for 2 minutes, 2% B for 2 minutes, 2–98% B for 2 minutes, 98% B for 2 minutes, 98–2% B for 2 minutes, and 2% B for 2 minutes. The orbitrap and FAIMS were operated in positive ion mode with a positive ion voltage of 1800V; with an ion transfer tube temperature of 270°C; using standard FAIMS resolution and compensation voltages of -50 and -65V, an inner electrode temperature of 100°C, and outer electrode temperature of 80°C with 4.6 ml/min carrier gas. Full scan spectra were acquired in profile mode at a resolution of 60,000, with a scan range of 400-1400 m/z, automatically determined maximum fill time, 300% AGC target, intensity threshold of 5 × 10^4^, 2-5 charge state, and dynamic exclusion of 30 seconds with a cycle time of 1.5 seconds between master scans. MS2 spectra were generated with a HCD collision energy of 32 at a resolution of 45,000 using TurboTMT settings with a first mass at 110 m/z, an isolation window of 0.7 m/z, 200% AGC target, and 120 ms injection time.

#### Phosphoproteomics time course analysis

Raw files were analyzed in Proteome Discoverer 2.4 (Thermo Fisher Scientific) to generate peak lists and protein and peptide IDs using Sequest HT (Thermo Fisher Scientific) and the ToxoDB release49 GT1 protein database. The unenriched sample search included the following post-translational modifications: dynamic oxidation (+15.995 Da; M), dynamic acetylation (+42.011 Da; N-terminus), static TMTpro (+304.207 Da; any N-terminus), static TMTpro (+304.207 Da; K), and static methylthio (+45.988 Da; C). The enriched sample search included the same post-translational modifications, but with the addition of dynamic phosphorylation (+79.966 Da; S, T, Y). The mass spectrometry proteomics data have been deposited to the ProteomeXchange Consortium via the PRIDE partner repository (Perez-Riverol et al., 2022) with the dataset identifier PXD033765 and 10.6019/PXD033765. Protein and peptide abundances are reported in **Supplementary File 1** and **Supplementary File 2**, respectively.

Exported peptide and protein abundance files from Proteome Discoverer 2.4 were loaded into R (version 4.0.4). Summed abundances were calculated for both DMSO and zaprinast-treated samples, incorporating the quantification values from both replicates. The ratio of zaprinast and DMSO peptide abundances was transformed into a modified Z-score. Peptides with modified Z-scores of 3 or above (839 peptides) or -1.8 or below (154 peptides) were considered dynamically changing. The normalized abundance values of peptides passing these thresholds were used as input for a principal component analysis using the R stats package (version 3.6.2). Clustering analysis was performed with the mclust package (Scrucca et al., 2016) (version 5.4.7) using the log_2_ ratios of zaprinast-treated samples relative to the DMSO-treated t = 0 samples. Cluster assignments are reported in **Supplementary File 3**.

#### PP_1_-AID phosphoproteomics

Samples were prepared as described above, with the following modifications. PP_1_-AID parasites were infected onto confluent HFF monolayers. IAA or vehicle (PBS) was added to the cell culture media 29 hours post-infection. At 32 hours post-infection, the monolayers were scraped, and parasites were mechanically released from host cells by passage through a 27-gauge needle. The parasites were harvested and concentrated as described above. The time-course was initiated when the parasite pellet was resuspended in 250 μl of 500 μM zaprinast solution. Aliquots of 50 μl were removed and combined with 2X lysis buffer at 0, 10, 30, and 60 seconds post-stimulation. The experiment was performed in biological duplicate, with vehicle- and IAA-treated parasites, for a total of 16 samples. The samples were prepared for mass spectrometry, labeled with TMTpro reagents, and enriched as described above. The mass spectrometry proteomics data have been deposited to the ProteomeXchange Consortium via the PRIDE partner repository (Perez-Riverol et al., 2022) with the dataset identifier PXD033765 and 10.6019/PXD033765.

Exported peptide and protein abundance files from Proteome Discoverer 2.4 were loaded into R (version 4.0.4). Peptides were clustered if they exhibited 2-fold difference in abundance between vehicle- and IAA-treated samples in both replicates for at least one time point. To focus on peptides dynamically changing during stimulation, we used for each peptide the grouped log_2_ ratios relative to the t = 0 abundance for each treatment as input to the mclust package (Scrucca et al., 2016) (version 5.4.7).

### Thermal profiling temperature range experiments

#### Parasite harvest and treatment

Temperature range thermal profiling experiments were performed in biological duplicate on different days. We have optimized thermal profiling for *T. gondii* and have published detailed descriptions of the protocol (Herneisen and Lourido, 2021). Confluent HFF cells in 15-cm dishes were infected with 2–5 ×10^7^ RH tachyzoites each. When the parasites had fully lysed the host cell monolayer (40-48 hours later), the extracellular parasites passed through a 5 μm filter. The parasite solution was concentrated by centrifugation for 10 minutes at 1000 x *g*. Parasites were resuspended in 1 ml of wash buffer (5 mM NaCl, 142 mM KCl, 1 mM MgCl_2_, 5.6 mM glucose, 25 mM HEPES pH 7.2) and spun again for 10 minutes at 1000 x *g*.

The parasite pellet was resuspended in 1200 μl of lysis buffer (5 mM NaCl, 142 mM KCl, 1 mM MgCl_2_, 5.6 mM glucose, 25 mM HEPES pH 7.2 with 0.8% IGEPAL CA-630, 1X Halt Protease Inhibitors, and 250 U/ml benzonase) and subjected to three freeze-thaw cycles. Half of the lysate was combined with an equivalent amount of 2X EGTA or 10 μM Ca^2+^ solution. The parasite extracts were incubated at 37°C/5% CO_2_ for 5 minutes. The parasite suspension was aliquoted into PCR tubes and was heated at the following temperatures on two 48-well heat blocks: 37, 41, 43, 47, 50, 53, 56, 59, 63, and 67°C. After 3 minutes, the tubes were removed from the heat blocks and were chilled on ice for 5 minutes. The lysates were transferred to a TLA-100 rotor and were spun at 100,000 *x g* for 20 minutes at 4 °C in a Beckman Ultra MAX benchtop ultracentrifuge. The solution, containing the soluble protein fraction, was removed for further processing.

#### Protein cleanup and digestion

The concentrations of the 37°C samples were determined with a DC Protein Assay (BioRad). A volume corresponding to 50 μg of the 37°C sample was used for further analysis, and equivalent volumes were used from the remaining temperature range samples. Samples were prepared using a modified version of the SP3 protocol (Hughes et al., 2019). 500 μg of a 1:1 mix of hydrophobic and hydrophilic beads (GE Healthcare 65152105050250 and 45152105050250) was added to each sample, followed by a 6X volume of 100% ethanol to induce aggregation. The samples were incubated at room temperature and 1,000 rpm on a thermomixer for 30 minutes. The beads were magnetically separated, and the supernatant was removed. The beads were washed three times with 80% ethanol followed by magnetic separation. Each sample was resuspended in 88 μl reduction buffer (10 mM TCEP and 50 mM TEAB, pH 8.5) and heated at 55°C for 1 hour. Samples were alkylated with 25 mM iodoacetamide shielded from light for 1 hour. 2 μg trypsin was added to each sample, and digestion proceeded overnight at 37°C on a thermomixer shaking at 1,000 rpm. The beads were magnetically separated, and the peptide eluatse were removed from further analysis.

#### TMT10plex labeling

Peptides were labeled according to the manufacturer’s protocol (Thermo Scientific 90111), with the following modifications. The eluates were combined with 200 μg of TMT10plex reagent in 41 μl of acetonitrile, for an estimated 1:4 w/w peptide:tag labeling reaction. The labeling proceeded for 1 hour at room temperature and was quenched for 15 minutes with 5% hydroxylamine. The samples were then pooled, flash-frozen, and lyophilized to dryness. The peptides were resuspended in 10% glacial acetic acid and were desalted using a Waters SepPak Light C18 cartridge according to the manufacturer’s instructions.

#### Fractionation

Samples were fractionated offline via reversed-phase high performance liquid chromatography. Samples were applied to a 10 cm × 2.1 mm column packed with 2.6 μm Aeris PEPTIDE XB-C18 media (Phenomenex) using 20 mM ammonium formate pH 10 in water as Buffer A, 100% acetonitrile as Buffer B, and Shimadzu LC-20AD pumps. The gradient was isocratic 1% Buffer A for 1 minute at 150 μl/min with increasing Buffer B concentrations to 16.7% B at 20.5 min, 30% B at 31 min and 45% B at 36 min. Fractions were collected with a FRC-10A fraction collector and were pooled to 8 samples per condition, followed by lyophilization.

#### MS data acquisition

The fractions were lyophilized and resuspended in 10-20 μl of 0.1% formic acid for MS analysis and were analyzed on a Q-Exactive HF-X Orbitrap mass spectrometer connected to an EASY-nLC chromatography system using 0.1% formic acid as Buffer A and 80% acetonitrile/0.1% formic acid as Buffer B. Peptides were separated at 300 nl/min on a gradient of 6–9% B for 3 minutes, 9–31% B for 100 minutes, 31–75% B for 20 minutes, and 75 to 100% B over 15 minutes.

The orbitrap was operated in positive ion mode. Full scan spectra were acquired in profile mode with a scan range of 375–1400 m/z, resolution of 120,000, maximum fill time of 50 ms, and AGC target of 3 × 10^6^ with a 15 s dynamic exclusion window. Precursors were isolated with a 0.8 m/z window and fragmented with a NCE of 32. The top 20 MS2 spectra were acquired over a scan range of 350–1500 m/z with a resolution of 45,000, AGC target of 8 × 10^3^, and maximum fill time of 100 ms, and first fixed mass of 100 m/z.

#### Thermal profiling temperature range analysis

Raw files were analyzed in Proteome Discoverer 2.4 (Thermo Fisher Scientific) to generate peak lists and protein and peptide IDs using Sequest HT (Thermo Fisher Scientific) and the ToxoDB release49 GT1 protein database. The search included the following post-translational modifications: dynamic phosphorylation (+79.966 Da; S, T, Y), dynamic oxidation (+15.995 Da; M), dynamic acetylation (+42.011 Da; N-terminus), static TMT6plex (+229.163 Da; any N-terminus), static TMT6plex (+229.163 Da; K), and static carbamidomethyl (+57.021 Da; C). Normalization was turned off. The mass spectrometry proteomics data have been deposited to the ProteomeXchange Consortium via the PRIDE partner repository (Perez-Riverol et al., 2022) with the dataset identifier PXD033713 and 10.6019/PXD033713. Protein abundances are reported in **Supplementary File 4**.

Protein abundances were loaded into the R environment (version 4.0.4) and were analyzed using the mineCETSA package (Dziekan et al., 2020), which performed normalization, generated log-logistic fits of the temperature profiles, and calculated a euclidean distance score (**Supplementary File 4**). Individual curve plots were generated using custom scripts, which are available upon request.

### Thermal profiling concentration range: Experiment 1

#### Parasite harvest and treatment

Concentration range thermal profiling experiments were performed in biological duplicate on different days. Confluent HFF cells in 15-cm dishes were infected with 2–5 × 10^7^ RH tachyzoites each. When the parasites had fully lysed the host cell monolayer (40-48 hours later), the extracellular parasites passed through a 5 μm filter. The parasite solution was concentrated by centrifugation for 10 minutes at 1000 x *g*. Parasites were resuspended in 1 ml of wash buffer (5 mM NaCl, 142 mM KCl, 1 mM MgCl_2_, 5.6 mM glucose, 25 mM HEPES pH 7.2) and spun again for 10 minutes at 1000 x *g*.

The parasite pellet was resuspended in 1200 μl of lysis buffer (5 mM NaCl, 142 mM KCl, 1 mM MgCl_2_, 5.6 mM glucose, 25 mM HEPES pH 7.2 with 0.8% IGEPAL CA-630, 1X Halt Protease Inhibitors, and 250 U/ml benzonase) and subjected to three freeze-thaw cycles. An equivalent volume of parasite lysate was combined with 2X [Ca^2+^]_free_ buffers to attain the final concentrations: 0 nM, 10 nM, 100 nM, 250 nM, 500 nM, 750 nM, 1 μM, 10 μM, 100 μM, and 1 mM. The solutions were aliquoted into two PCR tubes and were incubated for 5 minutes at 5% CO_2_/37°C. The tubes were placed on heat blocks pre-warmed at 54°C or 58°C for 3 minutes and were then immediately placed on ice. The lysates were transferred to a TLA-100 rotor and were spun at 100,000 *x g* for 20 minutes at 4 °C in a Beckman Ultra MAX benchtop ultracentrifuge. The solution, containing the soluble protein fraction, was removed for further processing.

#### MS Sample Preparation and Data Acquisition

Samples were prepared for mass spectrometry as described in the Temperature Range methods. MS data was acquired using the same instrument and methods as described above. Raw files were searched in Proteome Discoverer 2.4 using the search parameters described in the Temperature Range methods. Normalization was performed in the Proteome Discoverer software. The mass spectrometry proteomics data have been deposited to the ProteomeXchange Consortium via the PRIDE partner repository (Perez-Riverol et al., 2022) with the dataset identifier PXD033642 and 10.6019/PXD033642. Protein abundances are reported in **Supplementary File 5**.

### Thermal profiling concentration range: Experiment 2

#### Parasite harvest and treatment

Parasites and host cells were grown for one week in SILAC media and were considered separate biological replicates. SILAC media was prepared from DMEM for SILAC (Thermo Fisher 88364), 10% dialyzed FBS (Thermo Fisher A3382001) and L-Arginine HCl/L-Lysine-2-HCl (Thermo Fisher 89989 and 89987) or ^13^C_6_^15^N_4_ L-Arginine HCl/^13^C_6_ ^15^N_2_ L-Lysine-2HCl (Thermo Fisher 89990 and 88209). The parasites were harvested, lysed, and treated as described in the Concentration range: Experiment 1 procedure. The lysates were incubated on heat blocks pre-warmed at 50°C, 54°C or 58°C for 3 minutes and were then immediately placed on ice.

Insoluble aggregates were removed by filtration as described in (Herneisen and Lourido, 2021). In brief, the lysates were applied to a pre-equilibrated 96-well filter plate (Millipore MSHVN4510) and were spun at 500 *x g* for 5 minutes. The concentrations of the filtrates containing soluble fractions were quantified with a DC assay. The heavy and light samples were combined at 1:1 wt/wt, yielding an estimated 50 μg total per concentration.

#### Protein cleanup and digestion

Proteins were reduced with 5 mM TCEP for 20 minutes at 50°C. Alkylation of cysteines was performed with 15 mM MMTS for 10 minutes at room temperature. The samples were precipitated in 80% ethanol and were washed using the SP3 protocol as described in the Temperature range procedure. After the final wash, the samples were resuspended in 35 μl of digest buffer (50 mM TEAB) and 1 μg Trypsin). Digestion proceeded overnight at 37°C on a thermomixer shaking at 1,000 rpm. The beads were magnetically separated, and the peptide eluatse were removed from further analysis.

#### TMT10plex labeling

Peptides were labeled according to the manufacturer’s protocol (Thermo Scientific 90111), with the following modifications. The eluates were combined with 100 μg of TMT10plex reagent in 15 μl of acetonitrile, for an estimated 1:2 w/w peptide:tag labeling reaction. The labeling proceeded for 1 hour at room temperature and was quenched for 15 minutes with 5% hydroxylamine. The samples were then pooled, flash-frozen, and lyophilized to dryness.

#### Fractionation

Samples were fractionated with the Pierce High pH Reversed-Phase Peptide Fractionation Kit according to the manufacturer’s instructions for TMT-labeled peptides.

#### MS data acquisition

The fractions were lyophilized and resuspended in 10-20 μl of 0.1% formic acid for MS analysis and were analyzed on an Exploris 480 Orbitrap mass spectrometer equipped with a FAIMS Pro source (Bekker-Jensen et al., 2020) connected to an EASY-nLC chromatography system as described above. Peptides were separated at 300 nl/min on a gradient of 6–21% B for 41 minutes, 21–36% B for 20 minutes, 36–50% B for 10 minutes, and 50 to 100% B over 15 minutes. The orbitrap and FAIMS were operated in positive ion mode with a positive ion voltage of 1800V; with an ion transfer tube temperature of 270°C; using standard FAIMS resolution and compensation voltages of -50 and -65V (injection 1) or -40 and -60 (injection 2). Full scan spectra were acquired in profile mode at a resolution of 120,000, with a scan range of 350-1200 m/z, automatically determined maximum fill time, standard AGC target, intensity threshold of 5 × 10^3^, 2-5 charge state, and dynamic exclusion of 30 seconds with a cycle time of 2 seconds between master scans. MS2 spectra were generated with a HCD collision energy of 36 at a resolution of 30,000 using TurboTMT settings with a first mass at 110 m/z, an isolation window of 0.7 m/z, standard AGC target, and auto injection time.

#### Thermal profiling concentration range analysis

Raw files were analyzed in Proteome Discoverer 2.4 (Thermo Fisher Scientific) to generate peak lists and protein and peptide IDs using Sequest HT (Thermo Fisher Scientific) and the ToxoDB release49 GT1 protein database. The search included the following post-translational modifications for the first biological replicate, which was collected from parasites grown in light SILAC medium: dynamic phosphorylation (+79.966 Da; S, T, Y), dynamic oxidation (+15.995 Da; M), dynamic acetylation (+42.011 Da; N-terminus), static TMT6plex (+229.163 Da; any N-terminus), static TMT6plex (+229.163 Da; K), and static methylthio (+45.988 Da; C). The heavy samples were searched for the additional modifications Lys8-TMT6plex (+237.177 Da; K) and static Label:^13^C_6_^15^N_4_ (+10.008 Da; R). The mass spectrometry proteomics data have been deposited to the ProteomeXchange Consortium via the PRIDE partner repository (Perez-Riverol et al., 2022) with the dataset identifier PXD033650 and 10.6019/PXD033650. Protein abundances are reported in **Supplementary File 5**.

Protein abundances from Proteome Discoverer 2.4 were loaded into the R environment (version 4.0.4) and were analyzed using the mineCETSA package (Dziekan et al., 2020). The data were not further normalized using the package, as normalization had already been performed by the Proteome Discoverer software. Log-logistic fitting of the relative abundance profiles was performed using default settings. The AUC reflects the mean stability change when proteins were detected in both replicates of an experiment. If proteins were not detected, the AUC was calculated from one replicate. Proteins were considered Ca^2+^-responsive if the curve-fit parameters had an R^2^ > 0.8 and AUC two modified Z-scores from the median. The two concentration range experiments were analyzed separately. Individual curve plots were generated using custom scripts, which are available upon request.

### Enrichment Analysis

Sets of gene ontology terms from the differentially regulated (or Ca^2+^-responsive) and background proteome (all proteins with quantification values in each mass spectrometry experiment) were downloaded from ToxoDB.org (Molecular Function, Computed evidence, P-Value cutoff set to 1). Gene ontology terms were tested for enrichment across all gene ontology terms identified in the background proteome. A p-value for the likelihood of a given enrichment to have occurred by chance was obtained using a hypergeometric test.

### Immunoblotting

Samples were prepared as described in the thermal profile concentration range or phosphoproteomics procedures prior to proteomics sample preparation. The samples, which had already been treated with benzonase, were combined with 5X laemmli sample buffer (10% SDS, 50% glycerol, 300 mM Tris HCl pH 6.8, 0.05% bromophenol blue, 5% beta-mercaptoethanol) and were incubated at 37°C for 10 minutes. The samples were then run on precast 4–15% SDS gels (BioRad) and were transferred overnight onto nitrocellulose membranes at 4°C and 30 mA in 25 mM TrisHCl, 192 mM glycine, and 20% methanol. Blocking and antibody incubations were performed in 5% milk in TBS-T for 1 hour at room temperature. The membrane was washed three times with TBS-T between antibody incubations. Imaging was performed with the LICOR Odyssey CLx.

### Immunofluorescence assays

Confluent HFFs seeded onto coverslips were infected with extracellular parasites and were grown at 37°C/5% CO_2_. Approximately 21 hours later, IAA or a vehicle solution of PBS were added to the wells to a final concentration of 500 μM where indicated. At 24 hours post-infection, the media was aspirated, and coverslips were fixed in 4% formaldehyde in PBS. Following three washes in PBS, the fixed cells were permeabilized with 0.25% triton for 10 minutes at room temperature. Residual permeabilization solution was removed with three washes of PBS. The coverslips were incubated in blocking solution (5% IFS/5% NGS in PBS) for 10 minutes at room temperature, followed by a 60-minute incubation in primary antibody solution. An anti-CDPK1 antibody (Covance) was used as a parasite counterstain (Waldman et al., 2020). After three washes with PBS, the coverslips were incubated in blocking solution at room temperature for 5 minutes, followed by a 30-minute incubation in secondary antibody solution. The coverslips were washed three times in PBS and once in water.

Coverslips were mounted with Prolong Diamond and were set for 30 minutes at 37°C. Imaging was performed with the Nikon Ti Eclipse and NIS Elements software package.

### Invasion assays

Confluent HFFs seeded onto coverslips were incubated with 5 × 10^6^ extracellular parasites for 60 minutes at 37°C/5% CO_2_. The coverslips were washed four six times with PBS and were fixed for 10 minutes at room temperature with 4% formaldehyde in PBS. The coverslips were incubated in blocking solution (1% BSA in PBS) for 10 minutes. Extracellular parasites were stained with mouse anti-SAG1 for 30 minutes at room temperature. Following permeabilization with 0.25% triton-X100 for 10 minutes, the coverslips were incubated with guinea-pig anti-CDPK1 as a parasite counterstain for 30 minutes at room temperature. The coverslips were incubated with a secondary antibody solution containing Hoechst and were mounted on coverglass with Prolong Diamond. The number of parasites invaded was calculated by normalizing the number of intracellular, invaded parasites to host cell nuclei in a field of view. Five random fields of view were imaged per coverslip. Each experiment was performed in technical duplicate.

### Egress assays

Automated, plate-based egress assays were performed as previously described (Shortt and Lourido, 2020). In brief, HFF monolayers in a clear-bottomed 96-well plate were infected with 7.5 × 10^4^ or 1 × 10^5^ parasites of the TIR1 or PP_1_-AID strains, respectively. IAA or PBS were added to a final concentration of 500 μM 20 hours later. After 3 hours, the media was exchanged for FluoroBrite supplemented with 3% calf serum. Three images were taken before zaprinast (final concentration 500 μM) or A23187 (final concentration 8 μM) and DAPI (final concentration 5 ng /mL) were injected. Imaging of DAPI-stained host cell nuclei continued for 9 additional minutes before 1% Triton X-100 was injected into all wells to determine the total number of host cell nuclei. Imaging was performed at 37°C and 5% CO_2_ using a Biotek Cytation 3. Results are the mean of two wells per condition and are representative of three independent experiments.

### Replication assays

Parasites were inoculated onto coverslips containing HFFs. After 1 hour, the media was aspirated and replaced with media containing 500 μM IAA or PBS vehicle. At 24 h post-IAA addition, the coverslips with intracellular parasites were fixed, permeabilized, and stained with CDPK1 antibody and Hoechst as described under “Immunofluorescence assays”. For each sample, multiple fields of view were acquired with an Eclipse Ti microscope (Nikon) and the number of nuclei per vacuole were calculated from the full field of view (at least 100 vacuoles). Results are the mean of three independent experiments.

### Plaque assays

500 parasites were inoculated into 12-well plates of HFFs maintained in D10 and allowed to grow undisturbed for 7 days. IAA or vehicle (PBS) was added to a final concentration of 100 μM. Plates were washed with PBS and fixed for 10 min at room temperature with 100% ethanol. Staining was performed for 5 min at room temperature with crystal violet solution, followed by two washes with PBS, one wash with water, and drying.

### Live microscopy

PP_1_-mNG parasites were grown in HFFs in glass-bottom 35 mm dishes (Ibidi) for 24 hours. The media was decanted and the dish was washed once with 1 ml Ringer’s buffer (155 mM NaCl, 2 mM CaCl_2_, 3 mM KCl, 1mM MgCl_2_, 3 mM NaH_2_PO_4_, 10 mM HEPES, 10 mM glucose). Parasites were stimulated to egress with 500 μM zaprinast or 4 μM A23187 in Ringer’s buffer (155 mM NaCl, 2 mM CaCl_2_, 3 mM KCl, 1mM MgCl_2_, 3 mM NaH_2_PO_4_, 10 mM HEPES, 10 mM glucose) supplemented with 1% FBS (v/v) and recorded every 2 seconds for 300 seconds using an Eclipse Ti microscope (Nikon) with an enclosure maintained at 37 °C.

### Cytosolic Ca^2+^ measurements with FURA-2AM

Fura-2 AM loading of *T. gondii* tachyzoites was done as described previously (Moreno and Zhong, 1996). Briefly, freshly collected parasites (TIR1 or PP_1_-AID parasites treated with IAA for 5 hours) were washed twice with buffer A plus glucose or BAG (116 mM NaCL, 5.4 mM KCL, 0.8 mM MgSO_4_.7H_2_O, 50 mM Hepes pH 7.3, 5 mM Glucose) by centrifugation (706 x g for 10 min) and re-suspended to a final density of 1 x l0^9^ parasites/ml in loading buffer (BAG plus 1.5% sucrose, and 5 μM Fura-2 AM). The suspension was incubated for 26 min at 26 °C with mild agitation. Subsequently, parasites were washed twice by centrifugation (2000 x g for 2 min) with BAG to remove extracellular dye, re-suspended to a final density of 1×10^9^ parasites per ml in BAG and kept on ice. Parasites are viable for a few hours under these conditions. For fluorescence measurements, 2 × 10^7^ parasites/mL were placed in a cuvette with 2.5 mL of BAG. Fluorescence measurements were done in a Hitachi F-7100 fluorescence spectrometer using the Fura 2 conditions for excitation (340 and 380 nm) and emission (510 nm). The Fura-2 fluorescence response to Ca^2+^ was calibrated from the ratio of 340/380 nm fluorescence values after subtraction of background fluorescence at 340 and 380 nm as described previously (Grynkiewicz et al., 1985). Ca^2+^ release rate is the change in Ca^2+^ concentration during the initial 20 s after compound addition. Delta [Ca^2+^] was calculated by the difference between the higher Ca^2+^ peak and basal Ca^2+^.

## ACKNOWLEDGEMENTS

We thank Emily Shortt for assistance with cell culture, Eric Spooner of the Whitehead Proteomics Core facility for assistance with sample preparation, and Tyler Smith for providing assistance with tagging vector design. We thank L. David Sibley for the MIC2 and SAG1 antibodies, Dominique Soldati-Favre for the GAP45 antibody, and Marc-Jan Gubbels for the TUB1 antibody. This research was supported by funds from a National Institutes of Health grant to S.L. (R01AI144369) and a National Science Foundation Graduate Research Fellowship to A.L.H. (174530).

## FIGURES & LEGENDS

**Figure 1 Supplement.**
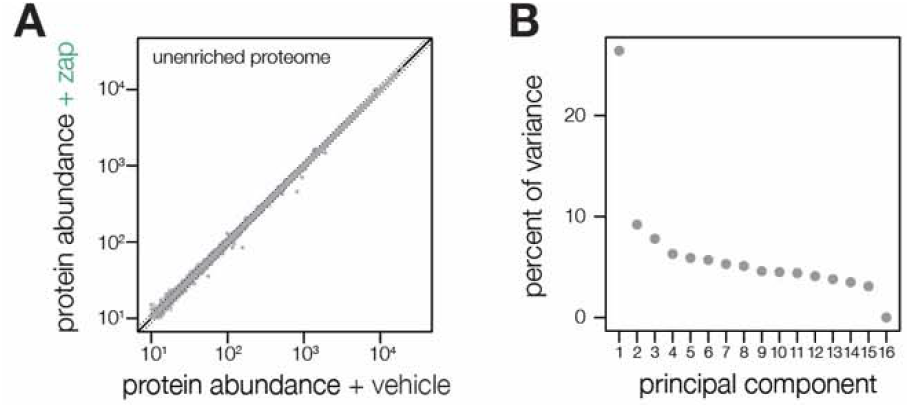
Metrics describing the zaprinast-dependent phosphoproteome. **(A)** Aggregate protein abundances for all time points from the non-phosphopeptide enriched samples of parasites treated with zaprinast or the corresponding vehicle (DMSO). Proteins quantified by a single peptide or more are shown in light and dark gray, respectively. Lines correspond to two median absolute deviations. **(B)** Proportion of the variance explained by each principal component, as described in **Figure 1C**.

**Figure 2 Supplement.**
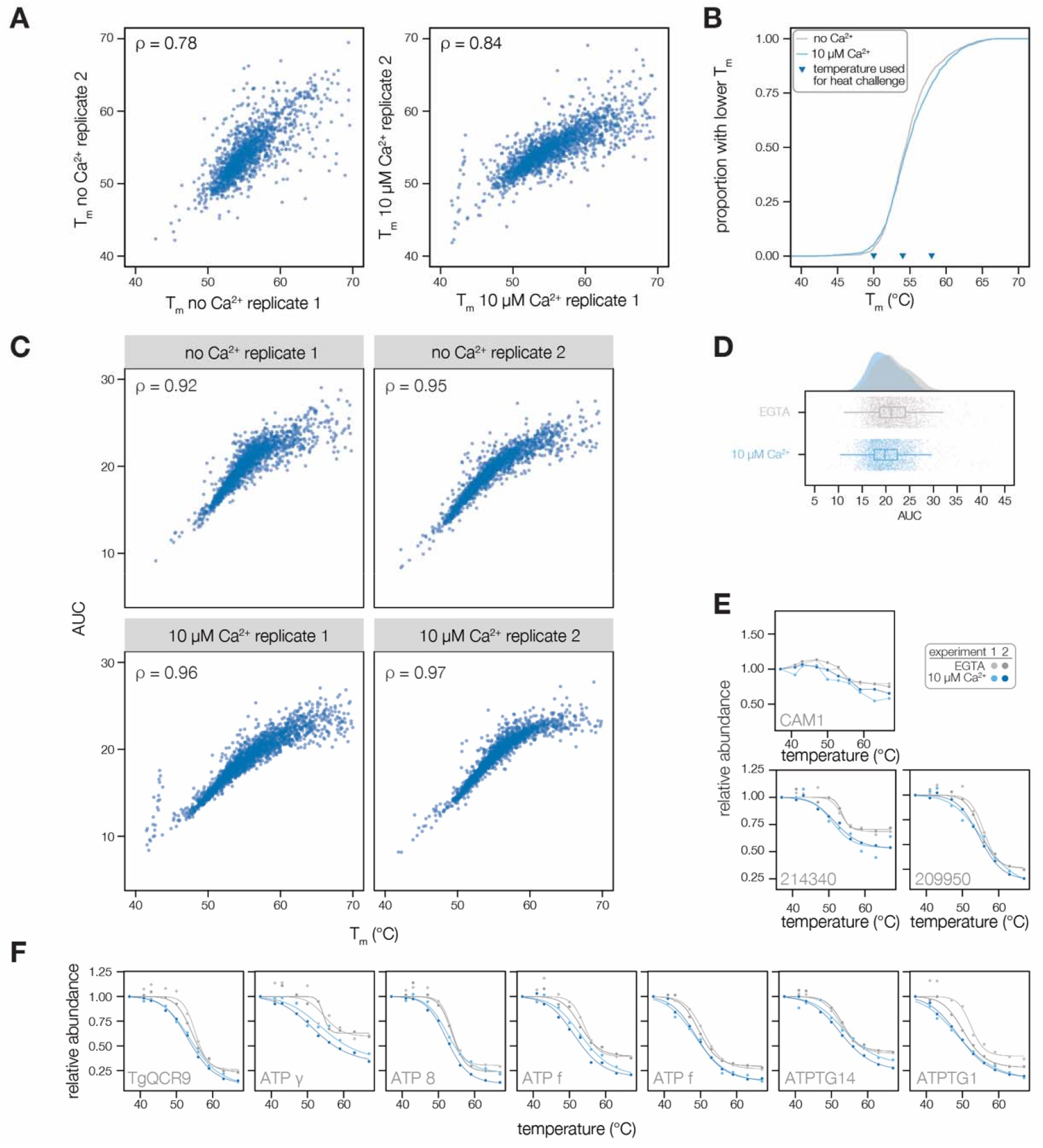
Extended data for thermal profiling experiments. **(A)** Comparison of the melting points of proteins with standard melting behavior (R^2^ > 0.8) across replicate experiments and treatment conditions. The correlation of melting temperatures is given by Spearman’s rho. **(B)** Cumulative distribution function of average measured melting temperatures in lysates with 10 μM or no Ca^2+^. **(C)** Correlation between T_m_ of proteins with standard melting behavior (R^2^ > 0.8) and AUC in each experiment. Correlation is given by Spearman’s rho. **(D)** Distribution of AUC (as in **Figure 1D**). **(E)** Melting curve of proteins discussed in the text: CAM1 (TGGT1_246930), an ICAP (TGGT1_214340, (Sidik et al., 2016a)), and a putative thioredoxin (TGGT1_209950). **(F)** Melting curves of *Tg*QCR9 and ATP synthase subunits (ATP) destabilized by Ca^2+^.

**Figure 3 Supplement.**
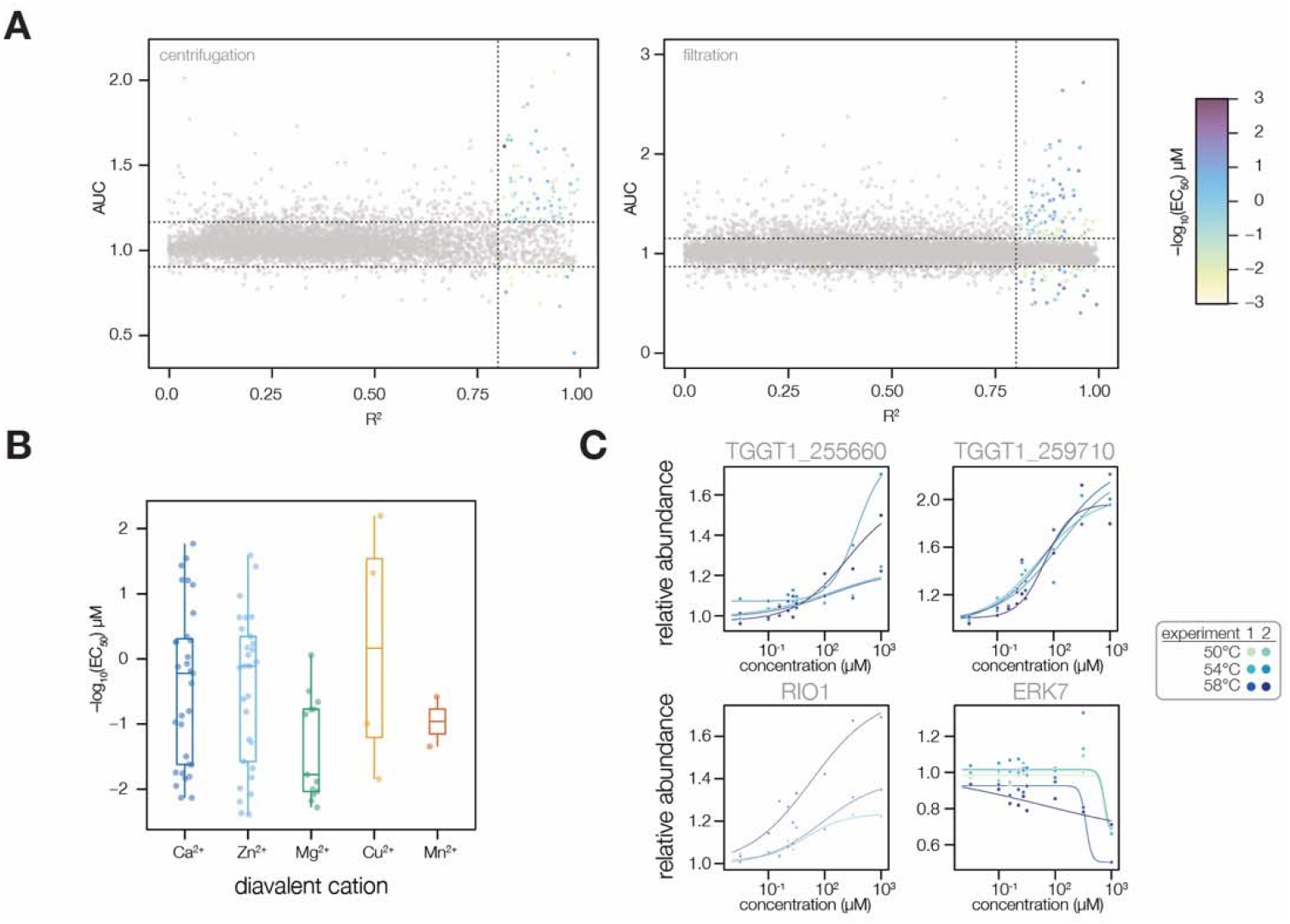
Extended analysis of thermal profiling experiments. **(A)** Plots of protein curve fit R^2^ vs. AUC, a measure of stability change, for each set of MS experiments. Dotted lines indicate thresholds for designated Ca^2+^-responsive behavior: R^2^ > 0.8 and an AUC two modified Z-scores from the median. Each point corresponds to an average of two replicates at each thermal challenge temperature (50, 54, or 58°C). Color denotes pEC_50_ in μM. **(B)** A comparison of the pEC_50_ values of proteins predicted to bind different divalent metal cations. Specificity was predicted via the presence of Interpro domains and through manual annotation. **(C)** Plots of individual protein melting curves, as described in the text: the EF hand domain-containing proteins TGGT1_255660 and TGGT1_259710; and the kinases RIO1 (TGGT1_210830) and ERK7 (TGGT1_233010).

**Figure 4 Supplement.**
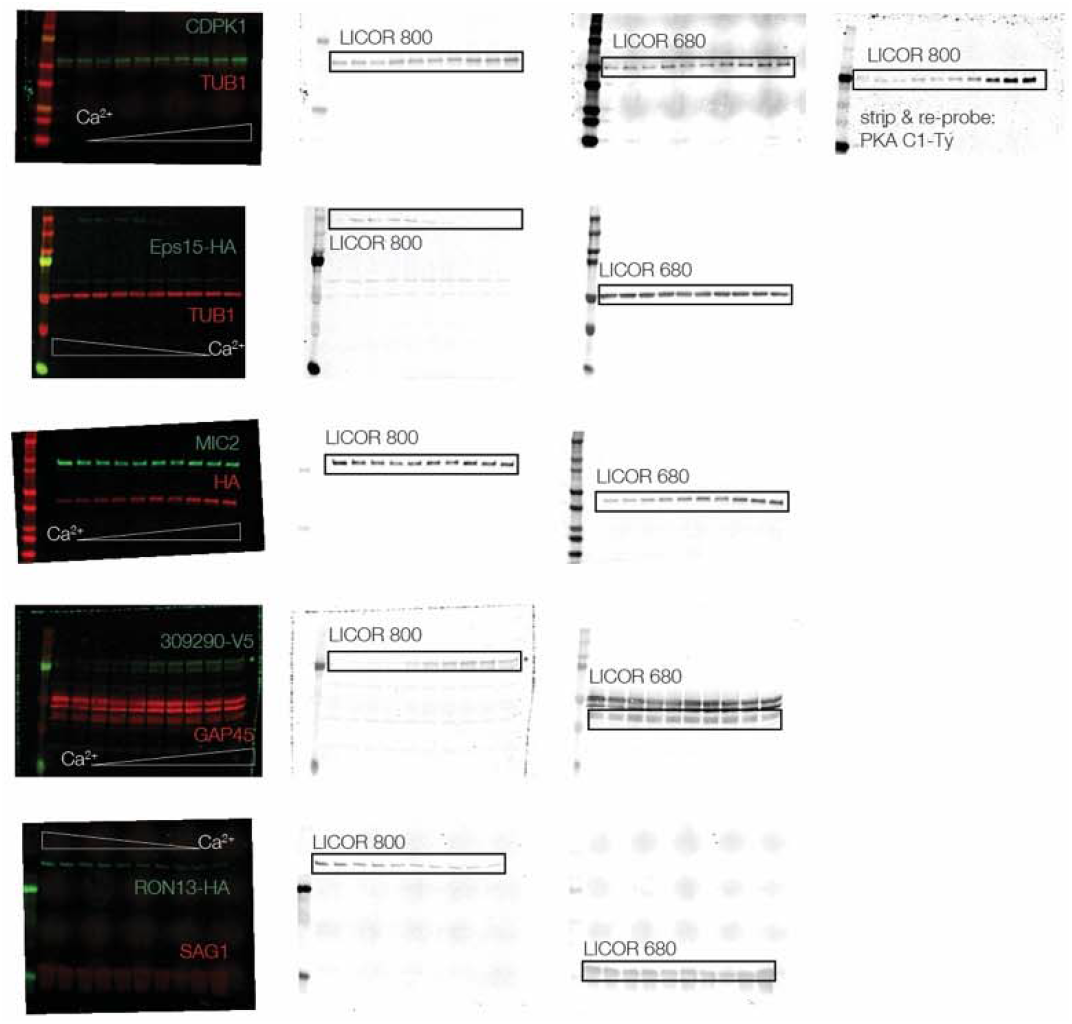
Uncropped immunoblots, as in Figure 4C. Parasite lysates were incubated at 10 Ca^2+^ concentrations and were thermally challenged at 58°C. Following centrifugation, the supernatant containing the soluble protein fraction was separated by SDS-PAGE. Following transfer onto a nitrocellulose membrane, the blots were probed with the indicated primary antibodies. The direction of the Ca^2+^ gradient is indicated on each blot. Representative blots from two biological replicates are shown.

**Figure 6 Supplement.**
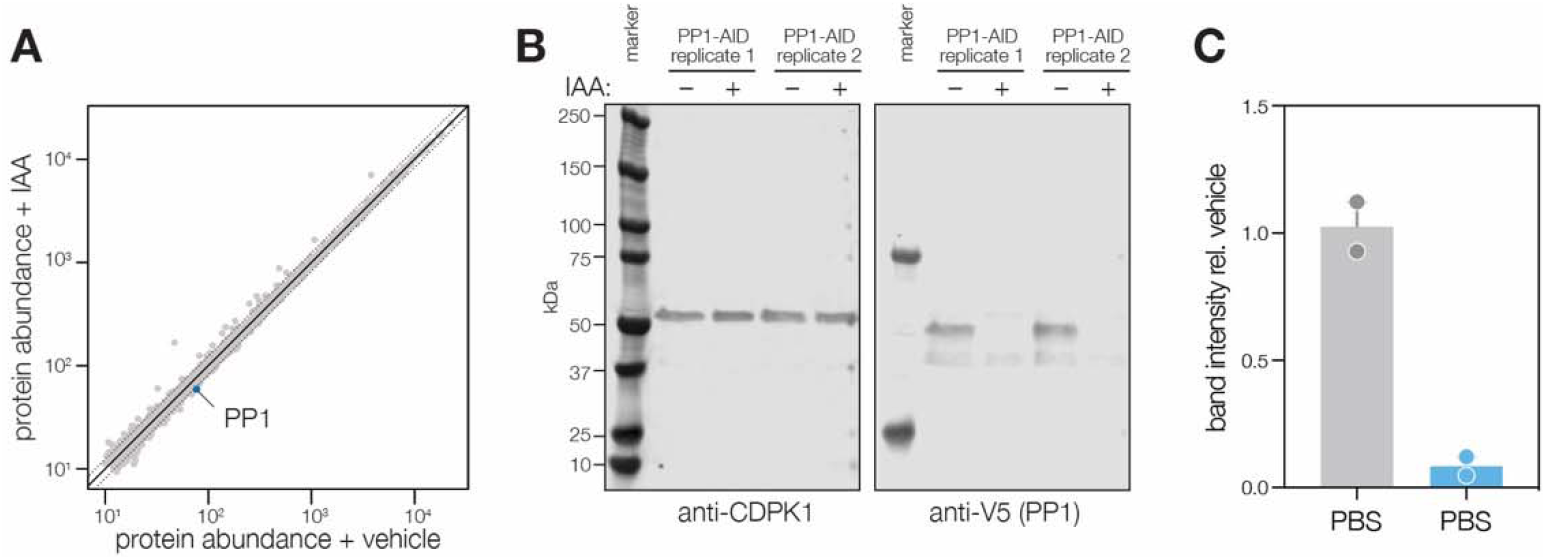
Extended analysis of PP_1_ phosphoproteomics experiment. **(A)** Aggregate protein abundances from the non-phosphopeptide enriched samples of parasites treated with IAA or vehicle. Proteins quantified by a single peptide or more are shown in light and dark gray, respectively. Lines correspond to two median absolute deviations. **(B)** Immunoblot of samples used for the PP_1_ phosphoproteomics experiment. **(C)** Quantification of immunoblot band intensity. Intensity was normalized relative to the signal of the vehicle-treated lane for each replicate.

**Figure 7 Supplement.**
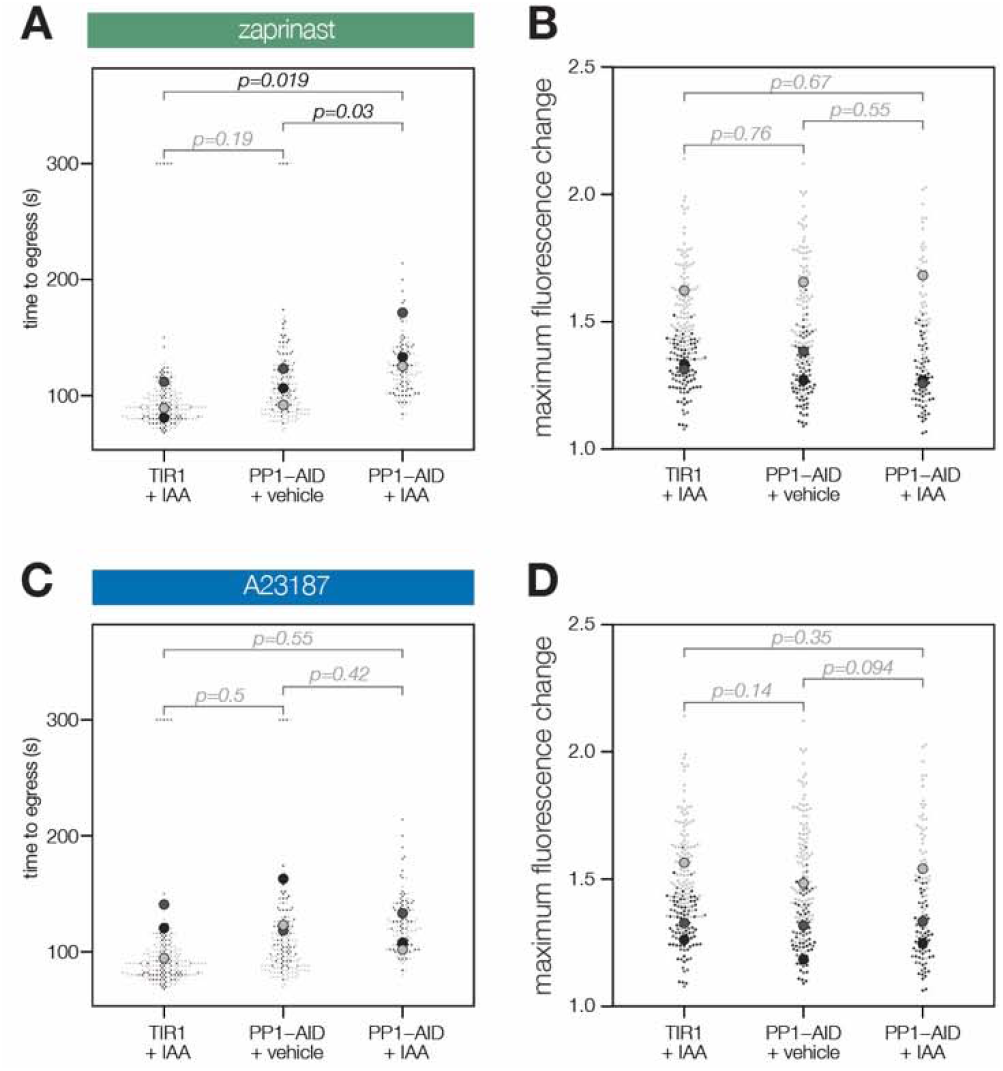
PP_1_-AID parasites egress, as quantified by video microscopy. **(A)** The time to vacuole egress after zaprinast treatment was manually scored. Different replicates are shown in different shades of gray. Transparent dots represent individual vacuoles; solid dots are the mean for each replicate. P-values were calculated from a two-tailed t-test. **(B)** The normalized fluorescence change of individual vacuoles after zaprinast treatment. Different replicates are shown in different shades of gray. Transparent dots represent individual vacuoles; solid dots are the mean for each replicate. P-values were calculated from a two-tailed t-test. **(C)** The time to vacuole egress after A23187 treatment was manually scored. Different replicates are shown in different shades of gray. Transparent dots represent individual vacuoles; solid dots are the mean for each replicate. P-values were calculated from a two-tailed t-test. **(D)** The normalized fluorescence change of individual vacuoles after A23187 treatment. Different replicates are shown in different shades of gray. Transparent dots represent individual vacuoles; solid dots are the mean for each replicate. P-values were calculated from a two-tailed t-test.

## SUPPLEMENTARY FILES

**Supplementary File 1**. Sub-minute phosphoproteomics time course protein and abundance assignments from Proteome Discoverer 2.4.

**Supplementary File 2**. Sub-minute phosphoproteomics time course peptide and abundance assignments from Proteome Discoverer 2.4.

**Supplementary File 3**. Mclust cluster assignments of phosphopeptides dynamically changing during zaprinast treatment.

**Supplementary File 4**. Data pertaining to the temperature range thermal profiling experiment.

Supplementary File 4.1. Protein and abundance assignments from Proteome Discoverer 2.4 for samples with 0 μM Ca^2+^, replicate 1.

Supplementary File 4.2. Protein and abundance assignments from Proteome Discoverer 2.4 for samples with 0 μM Ca^2+^, replicate 2.

Supplementary File 4.3. Protein and abundance assignments from Proteome Discoverer 2.4 for samples with 10 μM Ca^2+^, replicate 1.

Supplementary File 4.4. Protein and abundance assignments from Proteome Discoverer 2.4 for samples with 10 μM Ca^2+^, replicate 2.

Supplementary File 4.5. Curve fit output from the mineCETSA package.

Supplementary File 4.6. Area under the euclidean distance score calculations from the mineCETSA package.

**Supplementary File 5**. Data pertaining to the concentration range thermal profiling experiments.

Supplementary File 5.1. Protein and abundance assignments from Proteome Discoverer 2.4 for Experiment 1 samples with 54°C, replicate 1.

Supplementary File 5.2. Protein and abundance assignments from Proteome Discoverer 2.4 for Experiment 1 samples with 54°C, replicate 2.

Supplementary File 5.3. Protein and abundance assignments from Proteome Discoverer 2.4 for Experiment 1 samples with 58°C, replicate 1.

Supplementary File 5.4. Protein and abundance assignments from Proteome Discoverer 2.4 for Experiment 1 samples with 58°C, replicate 2.

Supplementary File 5.5. Curve fit output for concentration range Experiment 1 from the mineCETSA package.

Supplementary File 5.6. Area under the curve score calculations from the mineCETSA package for concentration range Experiment 2.

Supplementary File 5.7. Protein and abundance assignments from Proteome Discoverer 2.4 for Experiment 2 samples with 50°C, replicate 1.

Supplementary File 5.8. Protein and abundance assignments from Proteome Discoverer 2.4 for Experiment 2 samples with 50°C, replicate 2.

Supplementary File 5.8. Protein and abundance assignments from Proteome Discoverer 2.4 for Experiment 2 samples with 54°C, replicate 1.

Supplementary File 5.9. Protein and abundance assignments from Proteome Discoverer 2.4 for Experiment 2 samples with 54°C, replicate 2.

Supplementary File 5.10. Protein and abundance assignments from Proteome Discoverer 2.4 for Experiment 2 samples with 58°C, replicate 1.

Supplementary File 5.11. Protein and abundance assignments from Proteome Discoverer 2.4 for Experiment 2 samples with 58°C, replicate 2.

Supplementary File 5.12. Curve fit output for concentration range Experiment 2 from the mineCETSA package.

Supplementary File 5.13. Area under the curve score calculations from the mineCETSA package for concentration range Experiment 2.

**Supplementary File 6**. PP_1_ depletion zaprinast phosphoproteomics time course protein and abundance assignments from Proteome Discoverer 2.4.

**Supplementary File 7**. PP_1_ depletion zaprinast phosphoproteomics time course peptide and abundance assignments from Proteome Discoverer 2.4.

**Supplementary File 8**. Mclust cluster assignments of phosphopeptides dynamically changing during zaprinast treatment when PP_1_ is depleted.

